# Genomic Evidence for the Chinese Mountain Cat as a Wildcat Conspecific (*Felis silvestris bieti*) and Its Introgression to Domestic Cats

**DOI:** 10.1101/2020.12.07.414243

**Authors:** He Yu, Yue-Ting Xing, Hao Meng, Bing He, Wen-Jing Li, Xin-Zhang Qi, Jian-You Zhao, Yan Zhuang, Xiao Xu, Stephen J. O’Brien, Shu-Jin Luo

**Affiliations:** The State Key Laboratory of Protein and Plant Gene Research; Peking-Tsinghua Center for Life Sciences (CLS); Peking-Tsinghua-NIBS (PTN) Program; School of Life Sciences, Peking University, Beijing 100871, China; World Wide Fund for Nature, Beijing 100037, China; CAS Key Laboratory of Adaptation and Evolution of Plateau Biota and Qinghai-Tibetan Plateau Museum of Biology, Northwest Institute of Plateau Biology, Chinese Academy of Sciences, Xining, Qinghai 810008, China; Tibetan Plateau Wildlife Zoo, Xining, Qinghai 810001, China; Gansu Endangered Animals Protection Center, Wuwei, Gansu 733000, China; Laboratory of Genomic Diversity, Center for Computer Technologies, ITMO University, St. Petersburg 197101, Russia; Guy Harvey Oceanographic Center, Halmos College of Arts and Sciences, Nova Southeastern University, Ft Lauderdale, Florida 33004, USA

**Author notes:** Correspondence (S.J.L). Department of Archaeogenetics, Max Planck Institute for the Science of Human History, Jena 07745, Germany (He Yu).

## Abstract

The enigmatic Chinese mountain cat, endemic to the Qinghai-Tibet Plateau, has a controversial taxonomic status, whether a true species or conspecific with the wildcat *(Felis silvestris*) and whether it may have contributed to the domestication of cats *(F. s. catus*) in Asia. Here, we sampled 270 domestic and wild cats across China, sequenced 51 nuclear genomes, 55 mitogenomes, and multi-locus regions from modern and museum specimens. Genome-wide phylogenies supported taxonomic classification of the Chinese mountain cat as wildcat subspecies, *F. s. bieti*. No involvement of *F. s. bieti* in cat domestication in East Asia was detected, confirming that domestic cats shared a single origin from the African wildcat *(F. s. lybica*). A complex hybridization scenario including ancient introgression from the Asiatic wildcat *(F. s. ornata*) to *F. s. bieti*, and contemporary gene flow between *F. s. bieti* and sympatric domestic cats in the Tibetan region, raises the prospect of disrupting the genetic integrity of *F. s. bieti*, an issue with profound conservation implications.

## Introduction

The domestic cat *(Felis catus*, or *F. silvestris catus)* is one of the most popular pets in human society, with an estimated worldwide population over 600 million, including probably more than 100 million free-ranging feral cats *(1)*. Cat domestication origin and history have attracted wide public attention and scientific interest *(2)*. The first genetic study on the origin of domestic cats, based on mitochondrial and nuclear DNA assessment of nearly 1,000 specimens of domestic cats and their wildcat progenitors *F. silvestris*, revealed a single domestication event from the African wildcat *(F. s. lybica)* in the Near East *(3)*. These ancient cats probably coincided with the rise of early agriculture and civilization in the Fertile Crescent and subsequently expanded across the world. This conclusion was reinforced by an ancient DNA analysis of archaeological cat remains showing that African wildcats from both the Near East and Egypt contributed to the gene pool of modern domestic cats at different historical times *(4)*. Nevertheless, uncertainty remains whether multiple, independent cat domestication centers might exist, particularly given the lack of sampling in previous studies from the East Asia.

The wildcat, *F. silvestris*, from which domestic cats were derived, is widely distributed in the Old World and classified by controversial taxonomic systems, ranging from a monotypic taxon with multiple lineages to a species complex comprising at least two species *(5)*. According to the most recent genetic study which assembled wildcat samples worldwide *(3), F. silvestris* is resolved as a polytypic wild species including five distinct interfertile subspecies: *F. s. silvestris* the European wildcat, *F. s. lybica* from the Near East and northern Africa, *F. s. cafra* from southern Africa, *F. s. ornata* the Asiatic wildcat from central Asia east of the Caspian Sea, and *F. s. bieti* the Chinese mountain cat endemic to northwest China. By contrast, the Felidae taxonomy by Kitchener *et al*. *(5)* merged *F. s. cafra, F. s. lybica*, and *F. s. ornata* into *F. lybica* to unify wildcats from Africa to central Asia, meanwhile maintaining *F. silvestris* in Europe and *F. bieti* on the Qinghai-Tibet Plateau their own species status.

Two wildcat taxa, the Chinese mountain cat and Asiatic wildcat, are found in China. The Asiatic wildcat, *F. s. ornata*, occurs from the eastern Caspian Sea north to Kazakhstan, into western India, western China, and southern Mongolia, and its spotted coat pattern distinguishes it from other wildcat lineages that are typically striped. The Chinese mountain cat, *F. s. bieti*, also known as the Chinese desert cat or Chinese steppe cat, was initially described as an independent species *F. bieti* since 1892 *(6)*. It is the only wild felid endemic to China with a restricted distribution on the Qinghai-Tibet Plateau, and is characterized by a sand-colored fur with faint dark stripes, thick tail, light blue pupils, and ear tufts *(7)*. Molecular genetic studies suggested a reconsideration of Chinese mountain cat as conspecific of the wildcat based on its close association with other wildcat subspecies *(3, 8)*. This taxonomic revision is not yet unanimously accepted, as Kitchener *et al*. *(5, 9)* argued “*F. bieti* is morphologically distinct and is supposedly sympatric with *F. l. ornata*, which would also preclude its recognition as a subspecies of *F. silvestris/lybica*”. However, the presumed reproductive isolation between the Chinese mountain cat and Asiatic wildcat was based on their morphological divergence and possible overlapping distribution *(9)*, which, given the poorly defined range of the two taxa and possibly misidentification or mislabeling of specimens in previous studies, might not be supported.

Recent advances in genomic studies of exotic species have demonstrated that hybridizations between closely related taxa are common in nature and play an important role in shaping the genomes of modern animals *(10–14)*. Inter-taxa hybridization has also been documented in various lineages of Felidae, such as the big cats (genus *Panthera)* and Neotropical small cats (genus *Leopardus) (8, 15, 16)*. In northwest China, observations of cats possibly derived from interbreeding between the Chinese mountain cat and domestic cats are occasionally reported, leading to the postulation that local wildcats may have contributed to the gene pools of domestic cats in China. As one of the world’s oldest civilization centers, China is a known hotspot of animal and plant domestication, involving in or giving rise to numerous domesticated varieties such as the dog, pig, rice, and millet *(17–20)*. The earliest evidence of commensal relationship between human and a cat, in this case Asian leopard cat *(Prionailurus bengalensis)*, was also unearthed from a Neolithic site in northwest China *(21, 22)*, casting light on the existence of an environment conducive to a human-cat commensal process at that time in the East Asia.

On the other hand, genetic introgression from domestic species into their wild counterpart’s natural reservoirs is also evident in many taxa and may pose a threat to the wild populations by introducing deleterious traits or compromising their fitness in the wild *(23, 24)*. In some regions of Europe, the anthropogenic spread of domestic cats has resulted in range expansion of feral cats and the subsequent hybridization with the European wildcats *(25–29)*. Such widespread genetic infiltration from *F. s. catus* into *F. s. silvestris* has become a significant threat to the survival, distinctiveness, and genetic integrity of the sympatric European wildcat populations. On the Qinghai-Tibet Plateau where the Chinese mountain cat is endemic, most local domestic cats are free ranging, however, the circumstance and the extent of genetic admixture between domestic cats and wildcats remain elusive, let alone its potential conservation impacts on local wildlife.

To this end, we assembled thus far the most comprehensive set of samples of the Chinese mountain cat from its full range in the Tibetan region, the Asiatic wildcat from Xinjiang, and domestic cats across China especially from the regions sympatric and allopatric with the Chinese mountain cat, to resolve the phylogenetic mystery of one of the least studied felids in the world and to elucidate the evolutionary dynamics of the wildcats and domestic cats in East Asia. Data from whole genome sequencing and uniparental mtDNA and Y-chromosome molecular genetic markers jointly illuminated the Chinese mountain cat *F. s. bieti* and Asiatic wildcat *F. s. ornata* as equidistant and conspecific within the wildcat *(F. silvestris)*, an ancient introgression between *F. s. bieti* and *F. s. ornata*, and a complex pattern of contemporary gene flow from *F. s. bieti* into domestic cats across but not beyond its range. Domestic cats in China shared a Near Eastern origin with worldwide domestic cats, suggesting a single cat domestication origin from the African wildcat *(F. s. lybica)*.

## Results

### Range-wide sampling and initial genetic screening of wildcats and domestic cats

Sampling for this study covered a wide distribution of domestic cats in China and two of its wildcat relatives in the northwest China. The collection included four Asiatic wildcats *F. s. ornata* from Xinjiang and 27 Chinese mountain cats *F. s. bieti* across its full range on the Qinghai-Tibet Plateau, 12 of which were tissues or blood samples from road kills or zoo animals, and 15 pelts or bones from museums or local villages. In addition, blood, tissue, or saliva samples from 239 outbreed, unrelated domestic cats were gathered throughout China from 23 sites, including three locations within, three on the periphery of, and 17 distant from the core distribution of Chinese mountain cats *F. s. bieti* (Fig. 1A, Data file S1).

**Figure 1.**
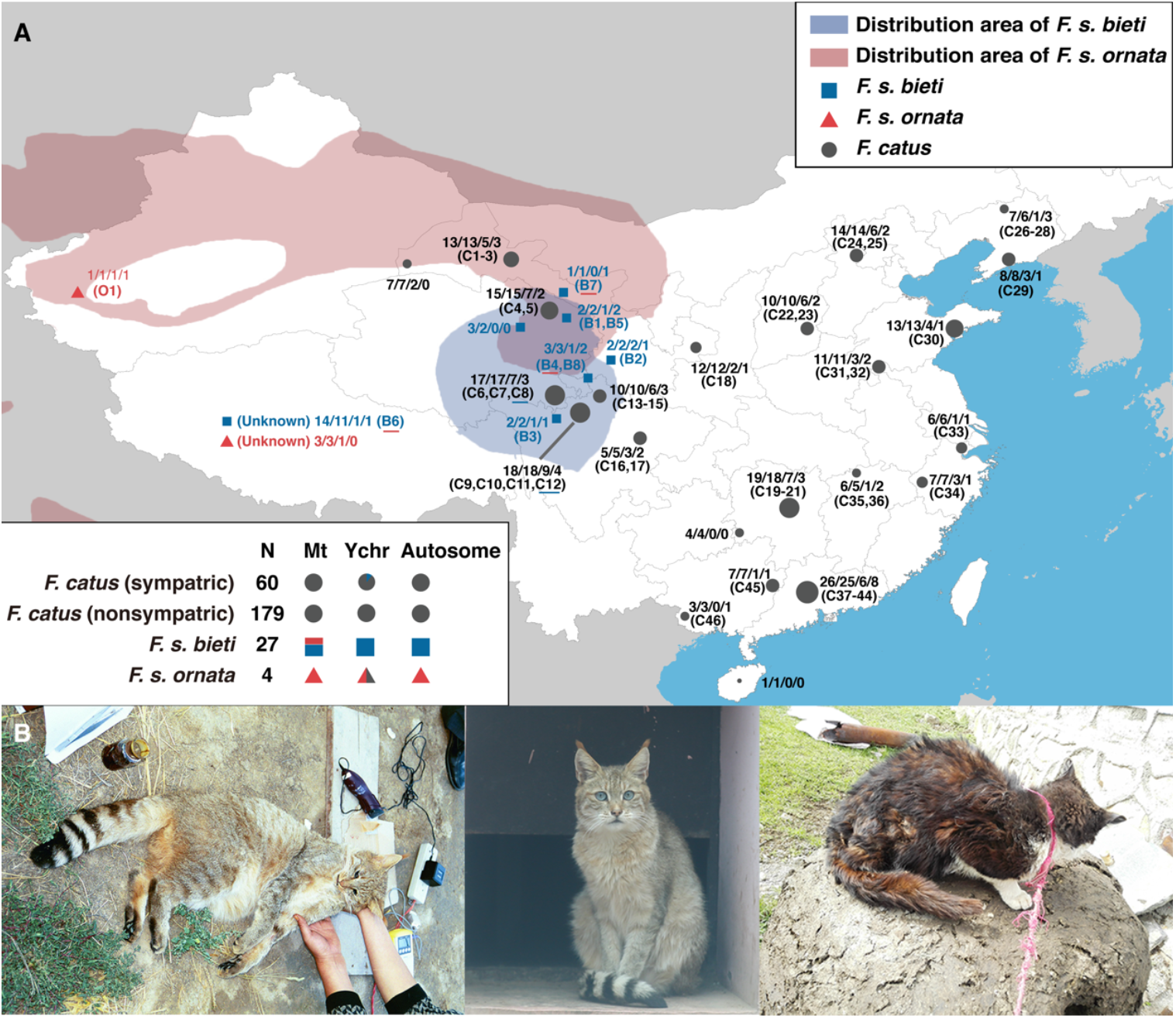
Wildcats *(Felis silvestris)* and domestic cats *(F. s. catus)* sampled from China in this study. **(A)** Sampling localities and range of wildcats and domestic cats in China. Labels at each sampling site indicate numbers of individuals following the order delimited by slash: (1) total sample size, with that of *F. s. catus* proportional to the area of circle; (2) samples with mtDNA fragment data; (3) samples with Y-chromosome fragment data; and (4) samples with WGS data and codes of animals were indicated below within parenthesis. Morphologically confirmed *F. s. bieti* individuals with *F. s. ornata* mtDNA haplotype and *F. s. catus* individuals with *F. s. bieti* Y-chromosome haplotype were underlined in red and blue on the map. The lower left panel summarizes sample sizes from each taxon and its population genetic background based on mtDNA, Y-chromosome and autosomal SNPs, with domestic cats from distribution area of *F. s. bieti* marked as “*F. s. catus* (sympatric)”, the others as “*F. s. catus* (nonsympatric)” and gray, blue and red colors corresponding to *F. s. catus, F. s. bieti* and *F. s. ornata* clades respectively. **(B)** Morphology of representative individuals from a “purebred” *F. s. bieti* (left), an *F. s. bieti* with *F. s. ornata* mtDNA haplotype (middle) and an *F. s. catus* with *F. s. bieti* Y-chromosome haplotype (right).

A multi-locus screening based on partial mtDNA and Y-chromosome sequencing was performed in all samples (261 specimens for mtDNA and 90 for Y-chromosome analysis) for an initial understanding of the genetic diversity pattern in the wildcat and domestic cats from China. Statistical parsimony network based on both markers revealed three distinct clusters that corresponded to the domestic cat *F. s. catus*, Asiatic wildcat *F. s. ornata*, and the Chinese mountain cat *F. s. bieti* (Fig. 2A). First, a 2,620 bp mtDNA fragment spanning *ND5, ND6*, and *CytB* were amplified in 250 modern specimens to distinguish 106 variable sites and 52 unique mtDNA haplotypes, including five haplotypes from 12 Chinese mountain cats *F. s. bieti*, three from four Asiatic wildcats *F. s. ornata*, and 44 from 234 domestic cats (Data file S2). One third (4 of 12) of the modern Chinese mountain cat specimens carried two mtDNA haplotypes that aligned with Asiatic wildcats, while the other eight individuals had three haplotypes exclusively found in the *F. s. bieti* clade. For degraded DNA extracted from museum samples, four short fragments within the 2.6 kb mtDNA haplotype were amplified separately and concatenated into 400-1,000 bp sequences (Table S1). Similar proportion of individuals with admixed genetic background was documented, as three of the 11 historical Chinese mountain cat specimens were different from the rest and contained variable sites diagnostic of the Asiatic wildcat (Data file S2).

**Figure 2.**
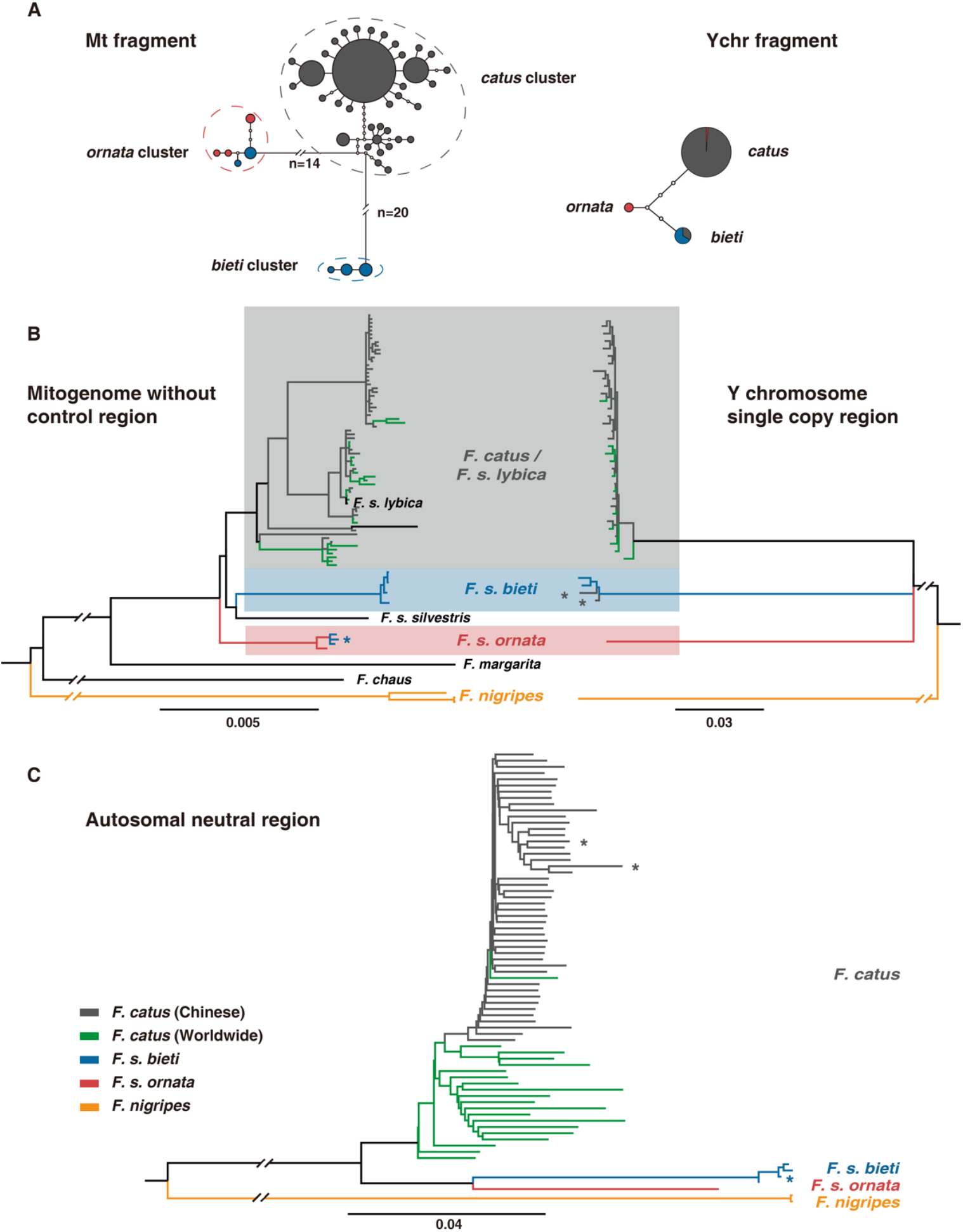
Phylogenetic relationships among wildcat lineages and domestic cats. **(A)** Statistical parsimony networks of *F. silvestris* and *F. s. catus* based on mtDNA and Y-chromosome fragments. The size of circles corresponds to number of individuals sharing this haplotype and the color corresponds to the taxonomic classification of the animal based on morphology (red - *F. s. ornata*; blue - *F. s. bieti*; gray - *F. s. catus)*. **(B)** Bayesian phylogeny of *Felis* spp. based on mitochondrial genome (excluding the control region) and Y-chromosome single-copy region. The branches are color-coded after the morphology-based taxonomic classification of the animal and the shaded boxes are color-coded corresponding to the three clades of interest in this study. The asterisks on certain branches mark the individuals with discordance between morphological appearance and genetic affiliation of certain genetic markers. **(C)** Phylogeny of *Felis* spp. based on genome-wide autosomal neutral SNPs reconstructed with neighbor joining method and distance matrix calculated following Gronau’s method. The branches are color-coded as in (B) and the asterisks on certain branches correspond to the same individuals marked in (B).

Two Y-chromosome fragments in *DBY7* and *SMCY7* were assembled from 90 male cats and three distinctive Y-haplotypes, each representing one of the three *Felis* taxa in the study, were identified based on six indels and SNVs from the concatenated 1,015 bp sequences (Table S1, Data S3). Patrilineal introgression was observed between the wildcat lineages and domestic cats from the Y-chromosome haplotype network. Three domestic cats, from Qinghai and Sichuan within the core range of Chinese mountain cats, shared the unique Y-haplotype found in all Chinese mountain cats, and one Asiatic wildcat showed a domestic cat Y-chromosome haplotype (Fig. 2A).

### Genome-wide phylogeny and taxonomy of the Chinese mountain cat

Fifty-five representative samples with adequate DNA quality, including eight Chinese mountain cats, one Asiatic wildcat, and 46 domestic cats, were proceeded for Illumina pair-end sequencing. Mitogenome and whole genome resequencing (WGS) data were generated from 51 samples at 6.8-15.1× coverage per individual, including four Chinese mountain cats (B1, B2, B3, and B4). Only the mitochondrial genome was reconstructed for the other four Chinese mountain cats (B5, B6, B7, and B8) due to sample quality constraint (Data file S4). Raw sequencing reads from 20 domestic cats and two black-footed cats *(F. nigripes)* were downloaded from NCBI Sequence Read Archive. A final dataset of 73 nuclear genomes with an average 16× coverage was obtained for variant calling and 20,425,451 biallelic autosomal SNPs were identified after quality filtering and masking. We also retrieved 67 different mitogenomes from 77 individuals, including six (mitogenome-B1, B2/B5, B3/B8, B4, B6, B7) from eight Chinese mountain cat specimens, and combined them with seven published mtDNA sequences (one from *F. s. bieti)* of the genus *Felis (8)* for downstream analyses.

Phylogenies inferred from mitogenome, Y chromosome, and the neutral region of autosomes, further illuminated the evolution and taxonomic classification of the Chinese mountain cat in relation with other wildcats and domestic cats. Phylogenetic inference based on mitochondrial sequences, exclusive of the control region, clusters all taxa, including the Asiatic wildcat *(F. s. ornata)*, European wildcat *(F. s. silvestris)*, Chinese mountain cat *(F. s. bieti)*, and a haplogroup consisting of all domestic cats *(F. s. catus)* and African wildcats *(F. s. lybica)*, into a single *F. silvestris* clade, with the Asiatic wildcat *F. s. ornata* situated as the basal lineage within the clade, though the support of the node was not strong (Fig. 2B; Fig. S1).

The patrilineal genealogy of 929 kb Y-chromosome single-copy region assembled from all available *Felis* spp. sequencing data also clustered *F. s. catus/F. s. lybica, F. s. bieti*, and *F. s. ornata* into one monophyletic group that is distinct from outgroup *F. nigripes*, despite an unresolved internal phylogeny among the three (Fig. 2B). In the neighbor-joining tree based on autosomal SNVs and average genomic divergence matrices *(30), F. s. bieti* (B1-B4) and *F. s. ornata* (O1) clustered as an early divergence prior to the domestic cat radiation among *F. silvestris* subspecies (Fig. 2C). Notably, all phylogenomic inferences placed the Chinese mountain cat within the *F. silvestris* subspecies clade, distinguished from other congeneric *Felis* species: the black-footed cat *F. nigripes*, the sand cat *F. margarita*, or the jungle cat *F. chaus*. The genome-wide phylogenetic patterns provided robust evidence for a close association of the Chinese mountain cat with the other *F. silvestris* taxa and corroborated the previously suggested reclassification of *F. bieti* as a subspecies of *F. silvestris (3, 8)*.

Domestic cats from China and other regions of the world were indistinguishable by either mtDNA or Y-chromosome phylogeny, in support of a single domestication event of worldwide cats from the African wildcat *(F. s. lybica)*. Nevertheless, all domestic cats from East Asia, including those from China and one from South Korea (W9, see Data file S4), formed a monophyletic group in the autosomal phylogeny, indicating a recent association among domestic cats from the region (Fig. 2C).

The discordance in the phylogenies inferred from maternal, paternal and bi-parental genetic markers are possibly the result of incomplete lineage sorting (ILS) and/or the existence of hybridization among lineages *(8)*. Consistent with the patterns from partial mtDNA and Y-chromosome genealogies, mitogenomes from three voucher Chinese mountain cats *F. s. bieti* (B4, B6, and B7, Fig. 1) clustered within the Asiatic wildcat *F. s. ornata*, and two domestic cats (C8 and C12, Fig. 1) carried Y-chromosome haplotypes embedded within the Chinese mountain cat clade (Fig. 2B). Intriguingly, genome-wide autosomal phylogeny (Fig. 2C) illustrated robust monophyly among *F. s. catus, F. s. bieti*, and *F. s. ornata* with no evidence of inter-lineage genetic admixture (thus excluding errors of morphological mis-identification). Bayesian coalescence analyses based upon mitogenome and Y-chromosome sequences both estimated the time to the most recent common ancestor (TMRCA) of *F. s. catus, F. s. bieti*, and *F. s. ornata*, at around 1.5 Mya during the Middle Pleistocene (Fig. S1), consistent with estimations from earlier studies *(8, 31)*. Such a relatively rapid and recent divergence of these lineages may have led to the phylogenetic discordance observed with different genealogies.

### Genetic introgression from the Chinese mountain cats to domestic cats

Principal component analysis (PCA) using autosomal neutral SNPs detected strong signal partitioning the three *F. silvestris* clades (Fig. 3A). The first principal component (PC1), which maximized 36% variance found among the individuals, distinguished black-footed cats from all *F. silvestris* taxa, indicative of a species-level divergence. The Chinese mountain cats, Asiatic wildcat, and domestic cats were separated along the PC2 which explained 10% of the variance, suggesting a divergence at the subspecific level. The PC3 revealed the intra-clade genomic diversity within domestic cats, segregating domestic cats of China from the worldwide cat populations. The alternative pairwise population genetic difference estimates also revealed a similar hierarchical partitioning of variances among the five groups (Table S2), with the F_ST_ between *F. nigripes* and the other four groups >0.7, while F_ST_ between *F. s. bieti, F. s. ornata* and *F. s. catus* was markedly lower (0.3 to 0.7), with F_ST_ between Chinese and worldwide domestic cat populations as low as 0.1.

**Figure 3.**
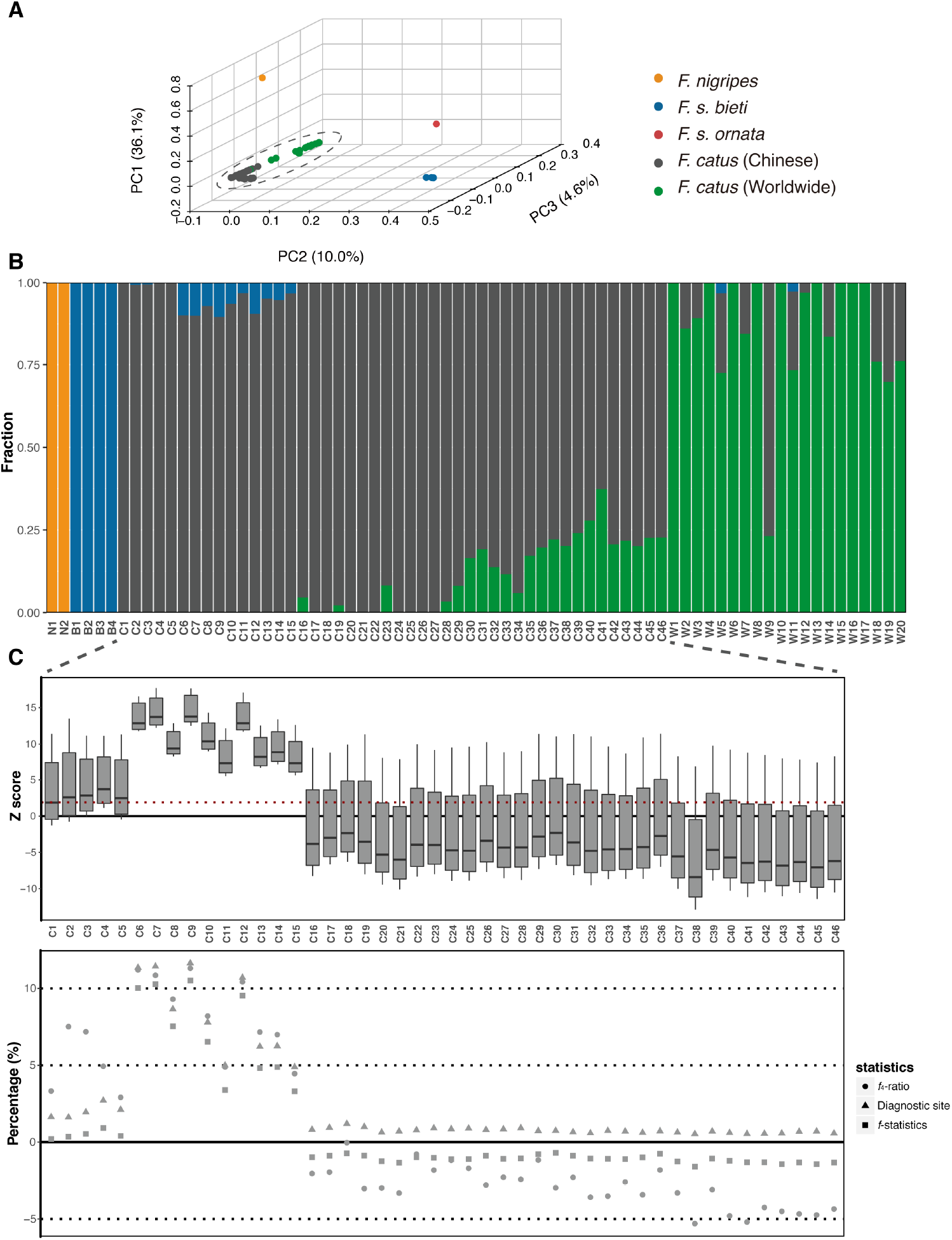
Population genetic structure and introgression from Chinese mountain cats to local domestic cats based on genome-wide neutral autosomal SNPs. **(A)** PCA of 73 individuals showing only the first three PCs. Two black-footed cats were separated from others with the first PC. *F. s. bieti, F. s. ornata* and *F. s. catus* were separated with the second PC. *F. s. catus* individuals were partitioned along the third PC with a moderate differentiation between domestic cats from China and worldwide. **(B)** Population structure of 72 individuals estimated in ADMIXTURE when K = 4. The four clusters correspond to *F. nigripes, F. s. bieti, F. s. catus* from China and *F. s. catus* worldwide, with ten *F. s. catus* (C6 - C15) carrying about 10% genetic admixture from *F. s. bieti*. **(C)** Genomic admixture between *F. s. bieti* and *F. s. catus* in China estimated by D-statistics, *f*_4_-ratio test, diagnostic sites and *f*-statistics. The D-statistics results were summarized with the boxplot showing distribution of Z scores with each *F. s. catus* in China and significant level (Z>2) showing with red dotted line. Percentage of diagnostic sites, *f*_4_-ratio and *f*-statistics were plotted to reveal the genomic admixture level from *F. s. bieti* of each *F. s. catus* in China.

A Bayesian analysis of 20 M SNPs as implemented in ADMIXTURE clustered 72 cats into four groups whose primary genomic affiliations correlated with black-footed cats, Chinese mountain cats, Chinese domestic cats, or worldwide domestic cats (Fig. 3B, *F. s. ornata* was excluded in the analysis due to limited sample size). It is worth noting though domestic cats sampled from the Chinese mountain cat’s core range in Sichuan and Qinghai (N=10, C6-C15 in Fig. 1A) carried around 10% genomic ancestry from *F. s. bieti*, indicating an extensive introgression from Chinese mountain cats to their sympatric domestic cats.

D-statistics was applied to further assess the extent of genetic admixture between Chinese mountain cats and domestic cats in China, using the worldwide domestic cats as a baseline. The level of wildcat’s genetic introgression in each domestic cat was quantified with fraction of diagnostic sites, *f*-statistics, and *f*_4_-ratio test (Fig. 3C, Table S3). All ten individuals sympatric with Chinese mountain cats presented significant signal of admixture in D-statistics, with the average Z-score ranging from 6.9 to 11.4, and the fraction of introgression between 4-12%, consistent with the estimated ancestry proportion in population clustering analysis (Fig. 3B). Intriguingly, a signal of introgression was also detected in five domestic cats (C1-C5) collected from northern Qinghai and Gansu, or the peripheral region of the Chinese mountain cats’ range (see Fig. 1A). The exact proportion of *F. s. bieti* introgression in these individuals varied between 0.5-7% with different methods, nevertheless significantly higher than that of domestic cats from areas beyond the geographic range of the Chinese mountain cat (Fig. 1A, Fig. 3C, Table S3). Overall, the genetic introgression from the Chinese mountain cats to domestic cats is restricted to the sympatric area in Qinghai, Sichuan, and Gansu, and the proportion of wildcat admixture in domestic cats decreases as the distance increases from the core to marginal distribution of the Chinese mountain cat. It is worth noting though none of the domestic cats with *F. s. bieti* genetic introgression can be distinguished from others by appearance, suggesting that morphological characters are not reliable diagnostic markers to identify hybrids.

The fine-scale distribution of the putative regions of introgression within the genomes of the admixed domestic cats (C6 - C15) was examined to determine whether the observed introgression in the sympatric domestic cats was introduced recently or ancient signals preserved in the population. Based on diagnostic SNPs, large continuous genomic segments over 30 Mb with Chinese mountain cat ancestry were identified in six of the ten domestic cats (Fig. S2). This indicated possible recent hybridization events between Chinese mountain cats and domestic cats as long-range linkage disequilibrium (LD) across the wildcat chromosomes had not been completely disrupted by recombination. The individual C8, a domestic cat from eastern Qinghai carrying a Chinese mountain cat-like Y-chromosome haplotype (Fig. 1B), displayed an extended homozygous region over 30 Mb with both alleles from Chinese mountain cat, consistent with a recent hybridization scenario reinforced by possible further inter-breeding between the fertile hybrid offspring.

The timing of the unidirectional introgression from the Chinese mountain cat *F. s. bieti* to its sympatric domestic cat in the Tibetan area was dated based on the extent of LD decay with Alder *(32)*. Domestic cats C6 - C15 from the core *F. s. bieti* range (namely “hybrid1”) and C1 - C5 from the boundary area of *F. s. bieti* distribution (namely “hybrid2”) were referred to as two admixed populations. The genomic introgression in hybrid1 population was well supported (p = 8.30E-16) with an exponential fit started at 2 cM (Table S4), and was estimated to have occurred around 7.42 generations ago (Fig. 4A). Using a generation time of two years for the domestic cat, the timing of the hybridization between Chinese mountain cats and domestic cats on the Qinghai-Tibet Plateau corresponded to 15 years ago. Similarly, we also detected significant signal of admixture (p = 8.90 E-05) in the hybrid2 population, which was estimated to be around 30.72 generations or 62 years ago (Fig. 4B).

**Figure 4.**
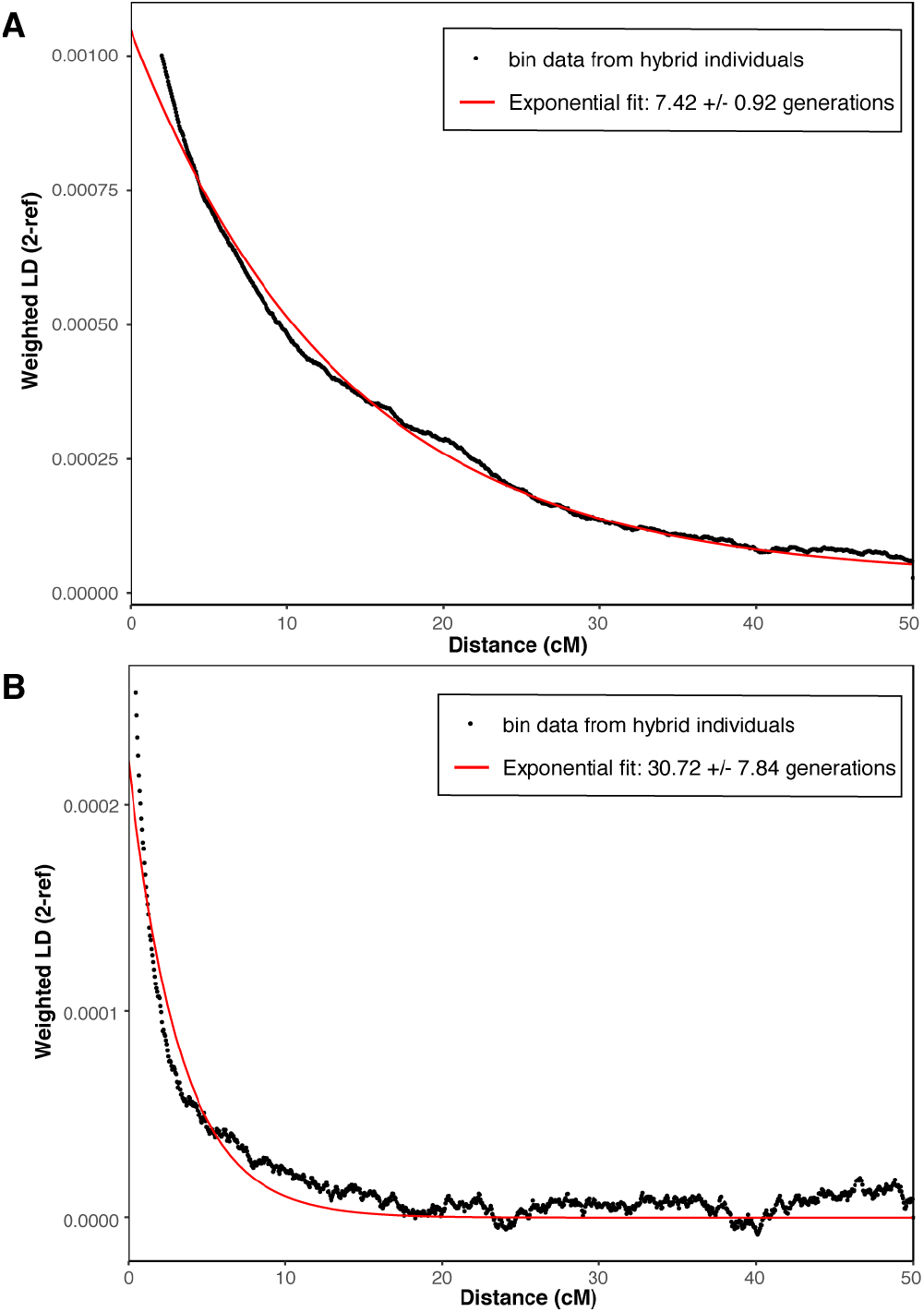
Dating of the hybridization between Chinese mountain cats and sympatric domestic cats based on LD decay. Weighted LD curves with putative pure Chinese domestic cats and Chinese mountain cats as two reference populations for **(A)** hybrid1 population with the exponential fit starting at 2.0 cM and **(B)** hybrid2 population the exponential fit starting at 0.5 cM.

### Evolutionary history of wildcats and domestic cats

The pairwise sequential Markovian coalescent (PSMC) model was employed to understand the demographic history, dispersal, and divergence in the wildcat and domestic cat clades within China (Fig. 5A). Both the Chinese mountain cat *F. s. bieti* and the Asiatic wildcat *F. s. ornata* displayed a moderate population expansion 1-2 million years ago (Mya) followed by a constant, gradual decline. The effective population size (Ne) of the African wildcat *(F. s. lybica)*, as represented by the genomic diversity of the domestic cat *(F. s. catus)*, experienced a significant rise around 100-400 thousand years ago (Kya) during the Middle to Late Pleistocene, which may reflect an ancient range expansion and/or population growth.

**Figure 5.**
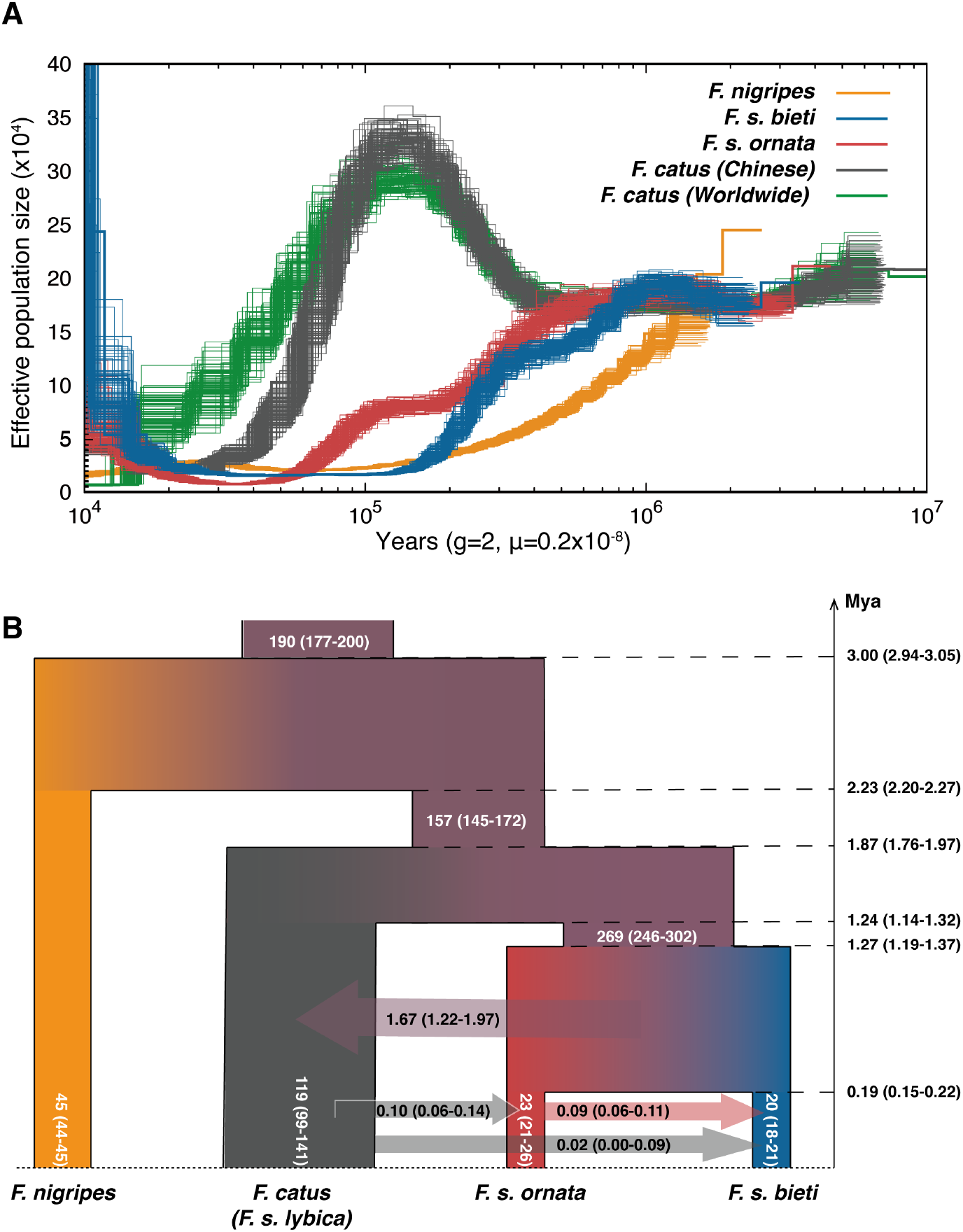
Demographic history of wildcats and domestic cats estimated by PSMC and G-PhoCS. **(A)** Population dynamics estimated by PSMC of five representative individuals from *F. nigripes, F. s. bieti, F. s. ornata, F. s. catus* from China and *F. s. catus* worldwide groups, with 100 bootstrap replicates. The results from *F. s. catus* individuals represent the dynamics of their wild ancestor, *F. s. lybica*. **(B)** Demographic model with divergence and migration among *F. nigripes, F. s. bieti, F. s. ornata* and *F. s. catus*, with the effective population sizes (in thousands), divergence times (in million years) and total migration rates estimated by G-PhoCS. The value and ranges in parenthesis represent the average and combined 95% Bayesian credible intervals of each estimated parameter and the branch width is proportional to effective population sizes.

The coalescent-based Generalized Phylogenetic Coalescent Sampler (G-PhoCS) was implemented to estimate the population divergence times and migration scenarios among *F. s. bieti, F. s. ornata*, and *F. s. catus/F. s. lybica*, using *F. nigripes* as the outgroup for time calibration (Fig. 5B). With a given topology as the phylogeny reconstructed with autosomal SNPs (Fig. 2C), 12 independent analyses were performed with all the combinations between one of the three domestic cats (C20, C25, W19) and one of the four Chinese mountain cats (B1-B4, Figure S3, Table S5). The coalescent time of *F. s. catus* and *F. s. bieti/F. s. ornata* lineages was estimated to be around 1.87 Mya (95% HPD at 1.76-1.97 Mya) and then *F. s. bieti* and *F. s. ornata* were coalescent to around 1.27 Mya (95% HPD at 1.19-1.37 Mya). We detected four significant inter-lineage migration bands, indicating the presence of gene flow from *F. s. catus* to *F. s. ornata* and from *F. s. ornata* to *F. s. bieti* with a total migration rates around 0.1, and from *F. s. bieti/F. s. ornata* lineage to *F. s. catus* with a total migration rate around 1.5 (Fig. 5B). The total migration rate from *F. s. catus* to *F. s. bieti* was minor and varied among different analyses from 0 to 0.09, with only one analysis including B2 showing significant level of gene flow. This observation confirmed that the hybridization between domestic cats and Chinese mountain cats was recent and hence the extent of genetic influence in *F. s. bieti* varied by individual. The effective population sizes estimated by G-PhoCS correlated well with PSMC, with the Ne of the ancestor of *F. s. catus/F. s. bieti/F. s. ornata* lineages around 157k, the Ne of the ancestor of *F. s. bieti* and *F. s. ornata* lineages increased to around 269k, and the current population sizes of *F. s. bieti* and *F. s. ornata* shrinking to about 20k and 23k respectively.

## Discussion

Whole genome sequencing data was retrieved from 46 domestic cats sampled across China. Both phylogenomic and population structure analyses clustered domestic cats from China and worldwide into one panmictic group (Fig. 2 and 3A, Table S2), thus supporting a single origin of domestic cats worldwide which is believed to have derived from the Near Eastern wildcat *(3, 4)*. However, autosomal phylogeny (Fig. 2C) and ADMIXTURE (Fig. 3B) also illuminated a close genetic association among domestic cats from China and South Korea that distinguished them from other populations in the world. This pattern implies a degree of isolation of domestic cats in East Asia after they dispersed or were introduced into this region. Although sampling of domestic cats in China specifically targeted local cats, while breed cats derived from regions outside China were avoided, genetic introgression from the worldwide cat population were detected in several individuals from southeastern China, which could be due to recent genetic interactions through introduction of breed cats from other countries into the gene pool of local feral cats.

There is no statistical evidence suggesting a significant contribution from local wildcats into the genetic ancestry of modern domestic cats in East Asia, yet a complex hybridization scenario among the contemporary domestic cat and wildcat lineages in the area was revealed. Genomic analyses indicate ancient admixture events between the Chinese mountain cat and the Asiatic wildcat, introgression from the domestic cat to the Asiatic wildcat (Figs. 2A, 5B), and recent genetic interactions between the Chinese mountain cat and its sympatric domestic counterparts (Figs. 2, 3) in the Qinghai-Tibet Plateau.

First, an ancient, unidirectional introgression from the Asiatic wildcat to the Chinese mountain cat was evident as only the “misplacement” of the Asiatic wildcat mtDNA haplotype was observed in Chinese mountain cats but not vice versa (Fig. 2), and only the migration band from *F. s. ornata* to *F. s. bieti* was significant in the G-PhoCS analysis (Fig. 5B). The signal of *F. s. ornata* admixture in *F. s. bieti* was only apparent in the maternally inherited mitochondrial lineages while genome-wide autosomal phylogeny and clustering algorithm supported monophyly for all morphologically distinguishable *F. s. bieti*. This cyto-nuclear discrepancy is consistent with an ancient admixture scenario in which mating of female *F. s. ornata* with male *F. s. bieti* occurred, followed by backcrossing of their offspring with *F. s. bieti* for a long period of time. Such asymmetric hybridization has been reported in various mammalian lineages, including Neotropical wild cats *Leopardus* spp., canids in North America, and African savannah and forest elephants *(15, 33–35)*, and was generally associated with population size contrast and mating preferences when two lineages encountered *(36)*. Likewise, the asymmetric introgression between the two wildcat lineages in Asia could be explained by the larger body size of the Chinese mountain cat relative to the Asiatic wildcat and hence a mating advantage in female choice selection of the latter. When the ancient *F. s. bieti* population overlapped with the range of the *F. s. ornata* whose population size was supposedly large, such admixture may occur and leave the signal in the genome of contemporary *F. s. bieti*.

Another unidirectional *F. s. lybica*/*F. s. catus*-to-*F. s. ornata* introgression was revealed via Y-chromosome genealogy and G-PhoCS analysis (Fig. 2A, 5B). This uni-directional inter-lineage gene flow can also be explained by difference in the population size of ancient African and Asiatic wildcat populations (Figure 5A) and/or the dispersal of post-domestication *F. s. catus* in Central Asia during the last millenniums. However, limited by the sampling size and the uncertainty of the exact geographic origin of the specimens, further study is necessary to examine whether the introgression occurred between the historical African and Asiatic wildcats prior to cat domestication, or between free-ranging domestic cats and Asiatic wildcats, a scenario resembling the genetic infiltration of feral domestic cats into the native European wildcats in Scotland *(37, 38)*.

As the only wildcat endemic to the Qinghai-Tibet Plateau of China, the genetic integrity of the Chinese mountain cat has been a subject of scientific interest and conservation concern. Genomic introgression from the Chinese mountain cat to its sympatric domestic cats was apparent and dating of the hybridization (Fig. 4) indicated it as a contemporary, not ancient, event. We observed a gradual decrease of *F. s. bieti* genetic constitution in domestic cats and older incidence of hybridization as they went from the center (e.g., Aba in western Sichuan and Golog in eastern Qinghai) to the margin (e.g., Jiuquan in eastern Gansu and Xining in northern Qinghai) of *F. s. bieti* range. The noticeable genetic admixture in domestic cat populations in the western Sichuan-eastern Qinghai boundary *(F. s. bieti* core range) and western Gansu-northern Qinghai area (peripheral range) were dated back to seven and 30 generations ago, corresponding to the beginning of the 21st century and mid-20th century, respectively. Since the signals of the earlier interbreeding could have likely been concealed by later events if multiple waves of population admixture recurred *(32)*, the contrast between the timing of admixture in domestic cats from different locations may reflect the continuous gene flow from Chinese mountain cats to domestic cats during the last century. It is also consistent with the pattern that more recent introgression is observed in the landscape where the Chinese mountain cat is abundant and in constant contact with domestic cats, whereas relatively older hybridization signals have been preserved in cats located in the peripheral region of *F. s. bieti* distribution.

Population census data from China recorded a dramatic increase of households and residents on the Qinghai-Tibet Plateau since the 1950s *(39)*, which coincided with the earliest *F. s. bieti* to *F. s. catus* admixture in Qinghai as documented in this study. Unlike the dog, the cat is not a domesticated animal generally associated with the traditional pastoral nomadic lifestyle of the Tibetan people and it is likely that the arrival and establishment of domestic cats on the Plateau was relatively recent. Regional socioeconomic development, migration of people from elsewhere into the highland, and the alteration in the local livelihood may have facilitated an expansion of free-ranging domestic cats, setting the stage for their close contact, frequent interaction, and possible interbreeding with the sympatric Chinese mountain cat. The exact population status of the Chinese mountain cat in the wild is unknown but nevertheless sparse and low-density *(40)*, therefore it may probably face a similar crisis like the European wildcat, that is, losing its genetic integrity and evolutionary adaptation to local environment to the introgression from an increasingly dominant local population of domestic cats *(26, 41)*.

Gene flow from the domestic cat to the Chinese mountain cat *F. s. bieti*, was detected in G-PhoCS analysis (Fig. 5B), despite a large variance in the estimates of total migration rates when different pairs of domestic cat and Chinese mountain cat were tested. Such fluctuation across individuals is consistent with recent introgression events, in which the extent of introgression varies by animals within the Chinese mountain cat population (Fig. S3). Unlike the abovementioned admixture analysis, no significant signal of gene flow was detected from Chinese mountain cats to domestic cats in G-PhoCS analysis. As the domestic cats used in G-PhoCS were from areas far from the the Chinese mountain cat range, this scenario can be explained by a contemporary admixture that was restricted to the local sympatric cats and exerted minor or no effect on domestic cat populations elsewhere.

The Felidae taxonomy by Kitchener *et al*. considers the Chinese mountain cat its own species while maintaining the other wildcat lineages as subspecies. In our population genomic analysis, the Chinese mountain cat, the Asiatic wildcat, and the domestic cat are equidistant, corroborating a subspecies-level recognition of these groups. Whole genome sequencing of the Chinese mountain cat, Asiatic wildcat, and domestic cats from China and worldwide, together with publicly available data from the European wildcat and African wildcat, provide firm support to the classification of the Chinese mountain cat as a subspecies of the wildcat, *F. s. bieti*. Phylogenetic analyses based on mitogenome, Y-chromosome, and genome-wide autosomal markers (Fig. 2) demonstrate a close monophyletic relationship between the Chinese mountain cat and other wildcat subspecies *(F. s. catus, F. s. ornata*, and *F. s. silvestris)* rather than a species-level distinctiveness *(42)*. The estimated timing of divergence between these *F. silvestris* subspecies is around 1.5 Mya, concordant to previous estimations based on nuclear sequence fragments and SNP array *(8, 31)*, significantly less than the divergence time between the accepted *Felis* species (i.e., *F. chaus, F. margarita, F. nigripes, and F. silvestris)* around 3 Mya. The PCA results also reflect a 3-fold less genetic distance among *F. silvestris* subspecies compared to their species-level divergence with *F. nigripes*.

Inter-lineage admixture between the Chinese mountain cat *F. s. bieti* and its closely related taxa is evident from this study, further advocating for an inclusion of all lineages as wildcat conspecific based on the biological species concept (BSC), which considers interbreeding as the prerequisite for a species *(43)*. A key argument from the proponents for the species status of the Chinese mountain cat lies on its distinctive morphological characters, a presumed sympatric distribution with the Asiatic wildcat, and an absence of gene flow between free ranging Chinese mountain cats and Asiatic wildcats *(9)*. However, recent surveys in Northwest China showed that the range attributed to the Asiatic wildcat may have been overestimated, as the presumed presence into plateau in northeastern Qinghai *(44)* may not hold true, thereby disputing the sympatric distribution of the two lineages. In addition, extensive genetic exchange between the two lineages is revealed as shown in the presence of “*F. s. ornata-*like” mitochondrial lineages in voucher Chinese mountain cats (Fig. 2). A significant migration band (total migration rate of 0.09) from the Asiatic wildcat to the Chinese mountain cat was also detected in demographic analysis with G-PhoCS (Fig. 5B). At last, the presence of interbreeding might diminish the morphological distinction as we have observed a Chinese mountain cat with “*F. s. ornata-*like” mtDNA haplotype does not show the typical thick and fluffy tail (Fig. 1B). Additional surveys and studies would help fine map the distribution of the Asiatic wildcat and Chinese mountain cat in Northwest China, to delineate the subspecies boundary or hybrid zone. Combined ecological and genetic data could elucidate the ancestry, adaptation, and evolution of these populations and resolve the historical and current patterns of gene flow among the wildcat and domestic cat lineages in the region.

## Conclusion

This study examined the genetic ancestry, population structure, and demographic history of wildcat and domestic cat lineages in East Asia from a whole genome perspective. Phylogenomic and population genomic analysis based on voucher specimens verified that the Chinese mountain cat *(F. bieti)*, a traditionally delineated felid species endemic to the eastern Qinghai-Tibet Plateau of China, is equidistant with other currently recognized wildcat lineages such as the Asiatic wildcat *(F. s. ornata)* and hence should be recognized as a conspecific, or *F. s. bieti*. Ancient introgression between *F. s. bieti* and *F. s. ornata* is evident as two deeply divergent mtDNA lineages were found within *F. s. bieti*. Domestic cats *(F. s. catus)* in China clustered with other cat populations worldwide, supporting a single Near Eastern origin of cat domestication from the African wildcat *(F. s. lybica)* and then spread globally. Contemporary genetic introgression from Chinese mountain cats into sympatric domestic cats is evident across but not beyond the range of *F. s. bieti*. The timing of admixture coincided with the large-scale socioeconomic change in the Tibetan area in the mid 20th century, which could have potentially led to an expansion of domestic cats into the region and is consistent with the scenario that domestic cats arrived rather late to the Plateau and thus have not encountered *F. s. bieti* until recently. The increasingly abundant local population of domestic cats may pose a threat to the Chinese mountain cat and jeopardize its genetic integrity and evolutionary adaptation to the high altitude, an issue with profound conservation implications and worth further studies.

## Materials and Methods

### Sample preparation

Samples were collected from 27 Chinese mountain cats, four Asiatic wildcats and 239 domestic cats, all with known geographic localities, to elucidate the evolutionary history of the sympatric *Felis* spp. in China. Specimens from the Chinese mountain cat included feces, blood, skin tissues, dry pelt, and skulls from zoos, museums, or local villages in Qinghai, Sichuan, and Gansu, and represented so far the largest range-wide sampling for this taxon (Data file S1). The source of the Asiatic wildcat samples in the study were blood or dry skin from southern Xinjiang. Buccal swab, blood, or skin tissues from outbred, unrelated domestic cats were collected from 23 sampling sites across China, with a particular emphasis on the areas sympatric and allopatric with the Chinese mountain cat (Fig. 1, Data file S1).

Genomic DNA from blood or skin tissues were extracted using a DNeasy Blood and Tissue Kit (Qiagen, Valencia, California, USA) and fecal samples were extracted using a QIAamp DNA Stool Mini Kit (Qiagen), following the manufacturer’s protocols. DNA from buccal swab samples collected using a PERFORMAgene PG-100 collection kit (DNA Genotek, Ottawa, Ontario, Canada) were extracted using buffer and protocol provided by the kit. DNA concentration and quality were examined with the NANODROP 2000 spectrophotometer and diluted to working solution for further analysis.

Genomic DNA extraction from museum samples were performed in a dedicated ancient DNA laboratory, following a modified silica-based spin column method and standard ancient DNA criteria with extreme precautions taken to minimize contamination risk from modern DNA samples and facilities *(45)*. For each specimen, 10-30 mg of skin tissues were pulverized in liquid nitrogen, washed twice with ddH_2_O and digested at 55°C overnight with 600 μL ATL buffer from DNeasy Blood and Tissue Kit (Qiagen), 24 mAu proteinase K (Qiagen), and 7 μL 1mol DTT. After digestion, DNA was purified using a silica column from QIAquick PCR purification kit (Qiagen) and kept at 4°C for downstream analysis.

### Multi-locus sequencing with mtDNA and Y-chromosome DNA markers

PCR primers were redesigned based on published *Felis silvestris* mtDNA sequences *(3)* to amplify a 2.7 kb mtDNA fragment spanning *ND5, ND6*, and *CytB*. Four short fragments ranging from 200-400 bp were selected within this region for the amplification of highly-degenerated DNA from museum samples. Two Y-chromosome DNA fragments encompassing the intronic regions of *DBY7* and *SMCY7* genes previously used in mammals and Felidae *(46, 47)* were amplified in all male individuals to examine their patrilineal ancestry.

The 2.7 kb mtDNA and Y-chromosome fragments were amplified in a 15 µL PCR reaction system containing 1X GC Buffer I, 1.0 mM dNTPs, 1 unit of TaKaRa LA Taq DNA polymerase (Takara Bio, Japan), 0.4 µM each of forward and reverse primers, and 10-20 ng of genomic DNA. For DNA extracted from museum specimens, PCR reactions were set up in the ancient DNA room following previously published protocol *(45)* and optimized for amplifying the short mtDNA fragments in a 25 µL PCR reaction system, which contained 1X PCR Buffer II, 5.0 mM MgCl_2_, 0.8 mM dNTPs, 10 µg BSA (Bovine serum albumin), 1 unit of AmpliTaq Gold DNA polymerase (Applied Biosystems, Waltham, Massachusetts, USA), 0.2 µM each of forward and reverse primers, and 5 µL of genomic DNA. PCR products were cleaned and sequenced on an ABI 3730XL sequencing system (Applied Biosystems) as described previously *(45)*. DNA sequences were inspected in Sequencher v5.0 (Gene Codes Co.) and concatenated into haplotypes for downstream analyses.

### Genome sequencing and NGS data processing

Illumina sequencing libraries with an insert size of 300-500 bp were constructed from 55 genomic DNA extracts following the manufacturer’s protocols (Illumina, San Diego, California, USA). The sample set consisted of eight Chinese mountain cats three of which carried the Asiatic wildcat mtDNA haplotypes, one Asiatic wildcat, and 46 local domestic cats across China two of which carried the Chinese mountain cat Y-chromosome haplotype (Fig. 1A). The libraries were sequenced on an Illumina HiSeq X Ten platform at Novogene Corporation to generate 150 bp paired-end reads. For four Chinese mountain cats with low DNA quality or endogenous DNA content, 2 Gb sequencing data per individual were produced for mitochondrial genome assembly. For the rest 51 samples including four Chinese mountain cats, one Asiatic wildcat, and all domestic cats, 30-40 Gb sequencing data per individual were generated for whole genome resequencing (WGS). In addition, WGS data of 20 domestic cats representing the worldwide population and two black-footed cats *(Felis nigripes)* were downloaded from the NCBI Sequence Read Archive (SRA) to be included in the analysis for comparison.

To exclude the interference from nuclear mitochondrial DNA segment (Numt) in assembling mitogenome, the sequencing reads from each individual were first mapped to the domestic cat mitogenome reference sequence (Accession: U20753) using Burrows-Wheeler Aligner (BWA) mem algorithm *(48)*. The mapped reads were then assembled into mitogenomes without the control region via a *de novo* genome assembly approach in Geneious v.9.1.5. From the 77 individuals sequenced from this or previous studies, 66 *Felis* spp. mitogenomes were assembled and identified for further analysis.

For genome-wide SNP identification and genotyping, the WGS reads of 73 individuals were mapped to the domestic cat genome assembly felCat8 and the domestic cat Y chromosome reference sequence (Accession: KP081775.1) using a BWA mem algorithm and default parameters. After removing PCR duplications and multi-targeted reads with SAMtools *(49)*, local realignment of the uniquely mapped reads were performed via RealignerTargetCreator and IndelRealigner in GATK v3.7 *(50)*. The reads realigned to autosomes and X chromosome were piled up using SAMtools for SNP calling in BCFtools *(51)*. The raw dataset of autosomal and X-chromosome SNPs was filtered for downstream analysis, which only retained those biallelic SNPs with Phred-scaled quality score over 20, raw read depth between 400 and 1600, genomic distance to the nearest indel over 5 bp, and no missing data across all the individuals. As phylogenetic and population genomic analyses were based on the putative neutral regions in the genome, we further excluded SNPs within the repetitive regions, CpG island regions, and protein-coding regions, based on the domestic cat felCat8 reference genome annotations from UCSC genome browser. The statistics of the WGS reads of each sample was summarized in Data file S4.

The realigned reads on Y chromosome from 40 males, including two Chinese mountain cats, 36 domestic cats, one Asiatic wildcat, and one black-footed cat, were piled up in SAMtools and genotypes were called as haploid using BCFtools. The initial dataset was filtered to keep only those biallelic SNPs with Phred-scaled quality score over 20, raw read depth between 100 and 400, and distance to the nearest indel over 5 bp. In order to eliminate the X-chromosome interference in the paternal genealogy analysis, Y-chromosome regions with X-homologue were identified by mapping sequencing reads from two females to the cat Y chromosome. After filtering only those SNPs located within the 929 kb single-copy Y-chromosome region *(52)* without female targeting were included in downstream Y-haplotype analysis.

### Phylogenomic analysis

Mitochondrial DNA and Y-chromosome haplotypes were aligned with Clustal X v2.0.10 *(53)* and the variable sites were identified with MEGA v6.06 *(54)* (Data file S2, 3). Statistical parsimony networks were constructed using TCS v1.1.3 *(55)* to infer the phylogenetic relationships among domestic cats and wildcats (Fig. 2A).

The 66 mitogenomes assembled from high-throughput sequencing data were aligned together with published mitogenome sequences from the domestic cat (Accession: U20753) and other *Felis* spp. (Accession: KP202273.1 - KP202278.1) for phylogeny reconstruction. The best fit nucleotide substitution model was selected using jModelTest v2.1.4. A Bayesian approach with two parallel Markov Chain Monte Carlo (MCMC) runs were performed in MrBayes v3.2.6 for 1,000,000 generations sampled every 500 generations. Phylogenetic analyses based upon maximum parsimony (MP), maximum likelihood (ML) with TrN (Tamura-Nei) +I+G model, and neighbor joining (NJ) constructed from Kimura two-parameter distances, were performed in PAUP v4.0b10, and the statistical reliability of each node was assessed by 100 bootstrap replicates.

The Y-chromosome phylogeny of 40 male cats was reconstructed following the same procedure but with HKY (Hasegawa-Kishino-Yano)+G model, and the ML analysis and bootstrap iterations were implemented in PhyML v3.1 *(56)*. The Bayesian trees based on mitochondrial genome and Y-chromosome were illustrated with Figtree v1.3.1, shown in Fig. 2B, and the bootstrap support or posterior probability of the tree topologies were marked in Fig. S1.

BEAST v2.4.4 *(57)* was applied to estimate the coalescence times of different wildcat lineages based on mtDNA and Y-chromosome SNPs. The mitogenome analysis was performed with TN93+I+G as the substitution model, the lognormal relaxed clock model, and the Yule tree prior. The coalescent times of *Felis* genus (4 Mya) and *F. silvestris* subspecies (1.5 Mya) were selected as two calibrations *(8, 31)*. The Y-chromosome analysis was performed with HKY+G as the substitution model, the lognormal relaxed clock model, and the Yule tree prior. The coalescent time between *F. nigripes* and *F. silvestris* (3 Mya), and subspecies of *F. silvestris* were used as calibrations *(8, 31)*. In both mitochondrial and Y-chromosome coalescence analyses, four parallel runs were performed for 50,000,000 generations with parameters sampled every 1,000 generations. Then log files and tree files were combined with the first 10% generations as burn-in to achieve the final estimation of parameters (Fig. S1).

To reconstruct the genome-wide autosomal phylogeny, a pairwise p-distance matrix based upon autosomal SNPs from the 73 individuals were computed according to Gronau *et al*. *(30)*. The neighbor-joining phylogenetic tree was constructed with the distance matrix using the Ape package v4.0 in R v3.2.3 *(58)*.

### Population genetic structure analysis

The principal component analysis (PCA) based on biallelic autosomal variants from all the individuals was performed with smartpca in EIGENSOFT v6.1.4 *(59, 60)* without removing outliers (Figure 3A). In addition, pairwise F_ST_ values among Chinese domestic cats, worldwide domestic cats, Chinese mountain cats, the Asiatic wildcat, and black-footed cats were estimated using VCFtools 0.1.15 *(61)* based on 1 Mb windows along the autosomes. Furthermore, population genetic structure of domestic cats and Chinese mountain cats was inferred from autosomal SNPs using ADMIXTURE *(62)* whereas the Asiatic wildcat was excluded from the analysis due to its extremely small sample size (N=1). The number of genetic clusters (K) was set from two to six and cross-validation statistics indicated K = 4 as the best fit for the dataset (Figure 3B).

### Gene flow detection and quantification

We applied D-statistics *(63)* as implemented in ADMIXTOOLS *(64)* to detect the presence of gene flow between Chinese mountain cats (termed “bieti”) and local domestic cats within China, with worldwide domestic cats (termed “worldwide”) for comparison and black-footed cats (termed “nigripes”) as the outgroup. All possible combinations of D (X, worldwide, bieti, nigripes) was calculated with X being each of the domestic cats from China and “worldwide” being each of the domestic cats from worldwide. The Z-score with each domestic cat from China was evaluated and those with the smallest Z-score over 2 were considered as domestic cats with genetic admixture from the Chinese mountain cat (Fig. 3C, Table S3).

Three statistical approaches were adapted to quantify the level of genetic introgression from Chinese mountain cats to these domestic cats, including calculating the percentage of Chinese mountain cat diagnostic sites in the domestic cat genomes, *f*-statistics *(65)*, and *f*_4_ ratio test *(66)*.

Autosomal SNPs specific to the Chinese mountain cat were identified by comparing the neutral genomic regions across four Chinese mountain cats to 20 worldwide domestic cats, whose chances of interbreeding with Chinese mountain cats were extremely low. The resultant 531,395 variants unique to the Chinese mountain cat were then used to calculate the portion of introgression in each of the admixed Chinese domestic cats.

The *f*_4_ ratio was calculated using ADMIXTOOLS with the following equation:

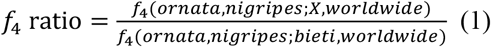

The *f*-statistics was calculated using ADMIXTOOLS with the D-statistics parameters following the equation below:

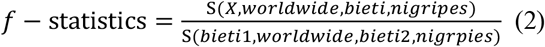

The numerator of D (X, worldwide, bieti, nigripes) (S-statistics) of each of the four Chinese mountain cats was divided by the numerator of D (bieti1, worldwide, bieti2, nigripes), in which two randomly selected Chinese mountain cats were designated as bieti1 and the other two as bieti2.

### Dating of the genetic introgression

The time of introgression from Chinese mountain cats to sympatric domestic cats was first estimated by plotting the genome-wide distribution of the Chinese mountain cat specific alleles in the 10 admixed domestic cats, along the 531,395 SNPs that distinguished the Chinese mountain cat and domestic cat (Fig. S2). Large consecutive genomic segments carrying the Chinese mountain cat ancestry within domestic cat genomes were then identified based on diagnostic variants. The time of the hybridization events was dated as implemented in Alder v1.03 *(32)* based on the patterns of linkage disequilibrium (LD) decay in two domestic cat populations, one including 10 cats from the core range of Chinese mountain cats (C6-15, “hybrid1”) and the other consisting of five individuals from the edge of the range (C1-5, “hybrid2”). One group with four Chinese mountain cats (B1-4) and the other with 31 domestic cats beyond the Chinese mountain cat distribution area (C16-46) were used as two ancestral populations in the analysis. To find the best fitting start point (d0), 11 parallel runs were conducted with d0 set from 0.5 cM to 5 cM, and 2.0 cM and 0.5 cM were selected as the best parameters for “hybrid1” and “hybrid2” populations according to p-values and Z-scores (Table S4).

### Demographic history inference

Pairwise Sequentially Markovian Coalescent (PSMC) model *(67)* and Generalized Phylogenetic Coalescent Sampler (G-PhoCS) *(30)* approach were applied to infer the demographic dynamics of wildcats and domestic cats in China, including historical population size, population divergence time, and gene flow scenarios.

PSMC was performed to estimate the change of effective population size through time, based on autosomal consensus sequences of representative individuals from five groups, namely N2, B4, O1, C25, and W19, which corresponded to *F. nigripes, F. s. bieti, F. s. ornata, F. s. catus* from China, and *F. s. catus* worldwide, respectively. The analysis was carried out at an individual-based level with 64 atom time intervals under the default pattern “4+25×2+4+6” as described by Li and Durbin *(67)* and the maximum coalescent time was set to 20. The estimated theta values were then transformed to effective population sizes and plotted with a generation time (g) as two years and the mutation rate (μ) as 2.4×10^−9^ substitution per site per generation (Fig. 5A). For each individual, 100 bootstrap replicates were run to evaluate the robustness of the estimation.

G-PhoCS was used to estimate the demographic parameters such as historical population size, divergence time, and migration rate based on coalescent-based Markov Chain Monte Carlo (MCMC) and a given topology *(30, 68, 69)*. In order to identify neutral loci for the analysis, the autosomal sequences of the hard-masked domestic cat genome assembly (felCat8) was further masked to remove CpG islands and exons with 1 kb franking regions based on UCSC genome annotations. Following the procedures described before *(30, 69)*, 34,418 unlinked loci were recognized, each of 1 kb in length, with a minimum inter-locus distance of 50 kb, and containing less than 10% masked sites.

G-PhoCS analysis was performed based on a given topology of the four *Felis* lineages and its estimated parameters (Figure S3A). Ten representative individuals were selected for the analysis, including four Chinese mountain cats (B1, B2, B3, B4), three domestic cats (C20, C25, W19), one Asiatic wildcat (O1), and one black-footed cat (N2). To avoid the possible interference between migration bands and time cost correlated with the complexity of demography model, we performed a prior analysis to identify significant migration bands with C25, B2, O1 and N2. All 18 possible migration bands were considered in the model and two parallel runs were conducted with 500,000 generations sampled every 100 generations, cross-checking all results to ensure convergence. Four significant migration bands were detected in this preliminary run with a total migration rate (mtot = m*tau) around or over 0.1 (Fig. S3B).

Then 12 independent analyses were run with four individuals from each of the four lineages and the four migration bands detected in the prior analysis, considering all the combinations of domestic cats and Chinese mountain cats. Each analysis was performed with two parallel runs, with 1,000,000 generations sampled every 100 generations and the trace files were combined with the first 30% generations as burn-in to obtain the final estimation of the demographic parameters (Figure S3C, Table S5). The estimated parameters (theta, tau) were converted to the effective population size (Ne), divergence time (T) and coalescent time (T_div) according to Gronau’s formulas *(30)*: theta = 4*Ne*μ, tau = T*μ/g and tau_div = tau+0.5*theta. The mutation rate was calibrated according to the divergence time between the black-footed cat and the domestic cat lineages (T_Felis_div) as 3 Mya *(8)*.

## Supplementary Materials

**Fig. S1**. TMRCA of wildcat and domestic cat lineages estimated from mitochondrial genome haplotypes and Y-chromosome SNPs.

**Fig. S2**. Genomic regions with introgression from *F. s. bieti* identified from ten sympatric domestic cats based on diagnostic SNPs.

**Fig. S3**. Topology model and parameters estimated by G-PhoCS with four *F. s. bieti*, three *F. s. catus*, one *F. s. ornata*, and one *F. nigripes*.

**Table S1**. Primers used in mitochondrial and Y-chromosome fragments amplification.

**Table S2**. Pairwise F_ST_ between different groups estimated based on 1Mb windows on autosomes

**Table S3**. Genetic introgression of *F. s. bieti* into *F. s. catus* from different areas of China.

**Table S4**. Alder fitting statistics with different fit starts estimated from weighted LD curves of two hybrid domestic cat populations.

**Table S5**. Summary statistics of G-PhoCS results, with average and combined 95% HPD of parameters from different runs and effective populations sizes, divergence times and total migration rates calculated with these estimated parameters.

**Data file S1**. Wildcat and domestic cat samples collected in the study, including the sample information and genetic information from preliminary analysis.

**Data file S2**. Haplotypes and variable sites identified in the 2.6 kb mtDNA fragment.

**Data file S3**. Haplotypes and variable sites identified in the 1016 bp concatenated Y-chromosome fragment.

**Data file S4**. Statistics of WGS reads generated and mapped to felCat8 assembly, domestic cat Y-chromosome and mitochondrial genome sequence.

## General

We thank LGD, NWIPB, Xining Zoo and Center of Endangered Animals in Gansu for providing wildcat samples and Chen Huaiqing, Daniel Gaillard, Huang Jianping, Xu Zinan, Zhang Pan, Wang Jinfan, Diao Ruxin, Li Peipei, Su Yaoxing, Li Zhong, Jin Bo, Zhang Houmei, Zhaxi Sange, Suori, Zhouba, Qiuguo, Liu Yongping,Wang Dehua, Bian Jianghui, Yang Qien, Carlos Driscoll, Nobuyuki Yamaguchi, and all the cat owners for their generous help in domestic cat sample collection. We are also grateful for the technique assistance and next generation sequencing service provided by Novogene.

## Author contributions

HY and SJL designed the experiment. HY, YTX and HM performed sample collection. HY finished the lab work and genomic analysis. HY and SJL wrote the manuscript. SJO helped revise the manuscript.

## Data and materials availability

All the newly generated sequencing data have been deposited in NCBI Sequence Read Archive project PRJNA478778.

## Figures and Tables

**Table 1.** Wildcat and domestic cat samples involved in the study.

| Taxonomy | Common name | Sample source | Total<br>No.<br>samples | MtDNA haplogroup |  |  |  | Y-chr haplogroup |  |  |  | No.<br>Samples<br>WGS |
| --- | --- | --- | --- | --- | --- | --- | --- | --- | --- | --- | --- | --- |
|  |  |  |  | bieti | ornata | catus | total | bieti | ornata | catus | total |  |
| *F. s. bieti* | Chinese | modern | 12 | 8 | 4 | 0 | 12 | 6 | 0 | 0 | 6 | 4 |
|  | mountain cat | museum | 15 | 8 | 3 | 0 | 11 | 0 | 0 | 0 | 0 | 0 |
|  |  | **Total** | **27** | **16** | **7** | **0** | **23** | **6** | **0** | **0** | **6** | 4 |
| *F. s. ornata* | Asiatic wildcat | **Total** | **4** | **0** | **4** | **0** | **4** | **0** | **1** | **1** | **2** | 1 |
| *F. s. catus* | Domestic cat | *F. s. bieti* core range<sup>1</sup> | 45 | 0 | 0 | 44 | 44 | 3 | 0 | 20 | 23 | 10 |
|  |  | *F. s. bieti* peripheral range<sup>2</sup> | 35 | 0 | 0 | 35 | 35 | 0 | 0 | 13 | 13 | 5 |
|  |  | Areas without *F. s. bieti*<sup>3</sup> | 159 | 0 | 0 | 155 | 155 | 0 | 0 | 47 | 47 | 31 |
|  |  | **Total** | **239** | **0** | **0** | **234** | **234** | **3** | **0** | **80** | **83** | **46** |

1. *F. s. bieti* core range refers to western Qinghai and northwestern Sichuan on the Qinghai-Tibet Plateau.
2. *F. s. bieti* peripheral range refers to the junction area of northern Qinghai and southern Gansu on the edge of the Qinghai-Tibet Plateau.
3. All other areas in China without any historical or present distribution of *F. s. bieti*.

