## Supplemental Figures and Tables for "Genomic Evidence for the Chinese Mountain Cat as a Wildcat Conspecific (*Felis silvestris bieti*) and Its Introgression to Domestic Cats"

**Supplementary Materials**

**Fig. S1.** TMRCA of wildcat and domestic cat lineages estimated from mitochondrial genome haplotypes and Y-chromosome SNPs. ……………………………………………………..2

**Fig. S2.** Genomic regions with introgression from *F. s. bieti* identified from ten sympatric domestic cats based on diagnostic SNPs. ……………………………………………………..4

**Fig. S3.** Topology model and parameters estimated by G-PhoCS with four *F. s. bieti*, three *F. s. catus*, one *F. s. ornata* and one *F. nigripes*. ………………………………………………..5

**Table S1.** Primers used in mitochondrial and Y-chromosome fragments amplification. …….7

**Table S2.** Pairwise $F_{ST}$ between different groups estimated based on 1Mb windows on autosomes. ………………………………………………………………………………………7

**Table S3.** Genetic introgression of *F. s. bieti* into *F. s. catus* from different areas of China. ..8

**Table S4.** Alder fitting statistics with different fit starts estimated from weighted LD curves of two hybrid domestic cat populations. ……………………………………………………...9

**Table S5.** Summary statistics of G-PhoCS results, with average and combined 95% HPD of parameters from different runs and effective populations sizes, divergence times and total migration rates calculated with these estimated parameters. ……………………….……10

**Data file S1-S4 are provided in Microsoft spreadsheet as different tabs.**

**Data file legends.** ..……………………………………………………………………….10

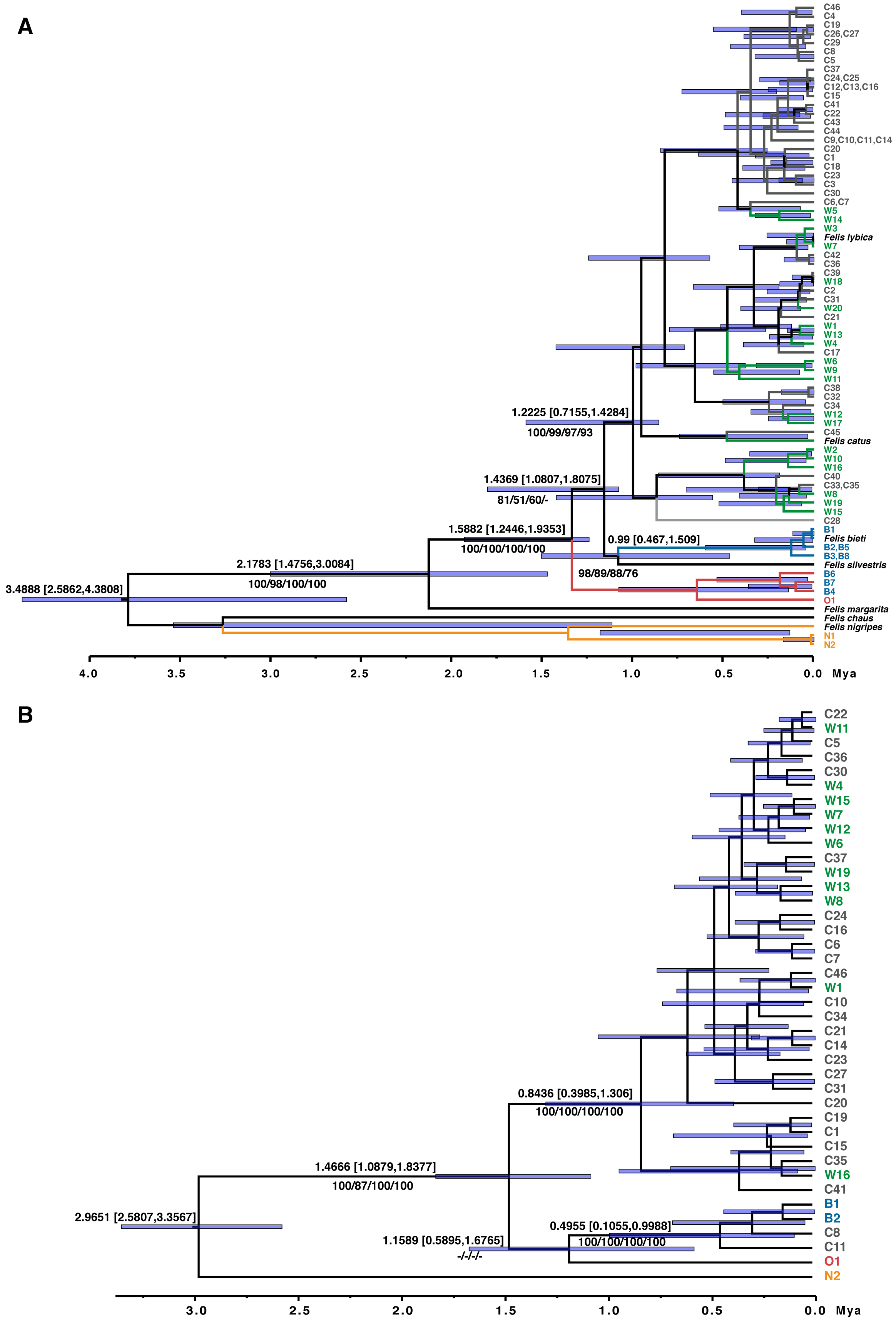
A
C46
C4
C19
C26,C27
C29
C8
C5
C37
C24,C25
C12,C13,C16
C15
C41
C22
C43
C44
C9,C10,C11,C14
C20
C1
C18
C23
C3
C30
C6,C7
W5
W14
W3
Felis lybica
W7
C42
C36
C39
W18
C2
C31
W20
C21
W1
W13
W4
C17
W6
W9
W11
C38
C32
C34
W12
W17
C45
Felis catus
W2
W10
W16
C40
C33,C35
W8
W19
W15
C28
B1
Felis bieti
B2,B5
B3,B8
Felis silvestris
B6
B7
B4
O1
Felis margarita
Felis chaus
Felis nigripes
N1
N2
1.2225 [0.7155,1.4284]
100/99/97/93
1.4369 [1.0807,1.8075]
81/51/60/-
1.5882 [1.2446,1.9353]
100/100/100/100
0.99 [0.467,1.509]
2.1783 [1.4756,3.0084]
100/98/100/100
98/89/88/76
3.4888 [2.5862,4.3808]
4.0
3.5
3.0
2.5
2.0
1.5
1.0
0.5
0.0 Mya
B
C22
W11
C5
C36
C30
W4
W15
W7
W12
W6
C37
W19
W13
W8
C24
C16
C6
C7
C46
W1
C10
C34
C21
C14
C23
C27
C31
C20
C19
C1
C15
C35
W16
C41
B1
B2
C8
C11
O1
N2
0.8436 [0.3985,1.306]
100/100/100/100
1.4666 [1.0879,1.8377]
100/87/100/100
2.9651 [2.5807,3.3567]
0.4955 [0.1055,0.9988]
100/100/100/100
1.1589 [0.5895,1.6765]
-/-/-/-
3.0
2.5
2.0
1.5
1.0
0.5
0.0 Mya

**Fig. S1.** TMRCA of wildcat and domestic cat lineages estimated from mitochondrial genome haplotypes and Y-chromosome SNPs.

(A) Time-calibrated tree of *Felis* genus estimated by BEAST2 with mitochondrial genomes without control region from this study and previous publications.

(B) Time-calibrated tree of wildcats and domestic cats estimated by BEAST2 with Y chromosome SNPs in SCR region identified in this study.

Numbers above the branches are the means and 95% HPD intervals of node ages estimated with Yule Model and numbers below the branches are support rates of nodes estimated with Bayesian/ML/MP/NJ methods. Color of the names indicates the taxonomy of individuals according to Figure 2C.

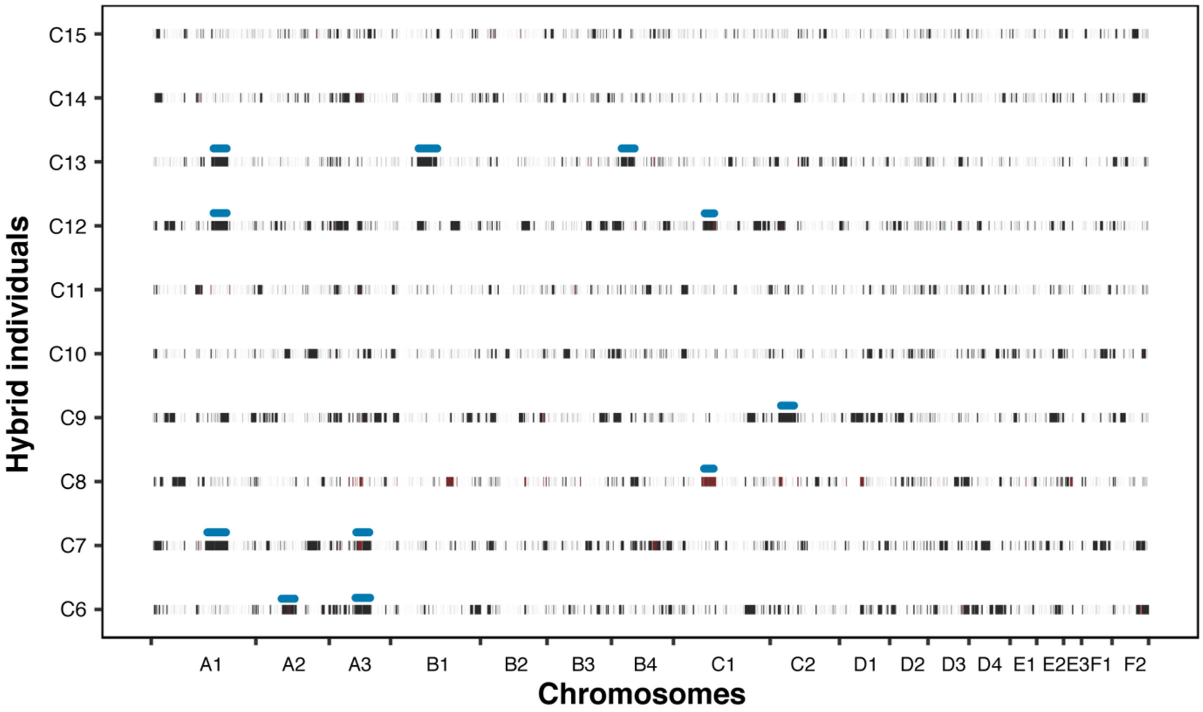


**Fig. S2.** Genomic regions with introgression from *F. s. bieti* identified from ten sympatric domestic cats based on diagnostic SNPs.

The black and red vertical lines indicate diagnostic alleles representing *F. s. bieti* ancestry in one haplotype or two haplotypes respectively. Chromosome names are listed at the X-axis and individual names listed at the Y-axis. The blue bars indicate chromosome segments of putative recent ancestry from *F. s. bieti* with their approximate length marked above the bars.

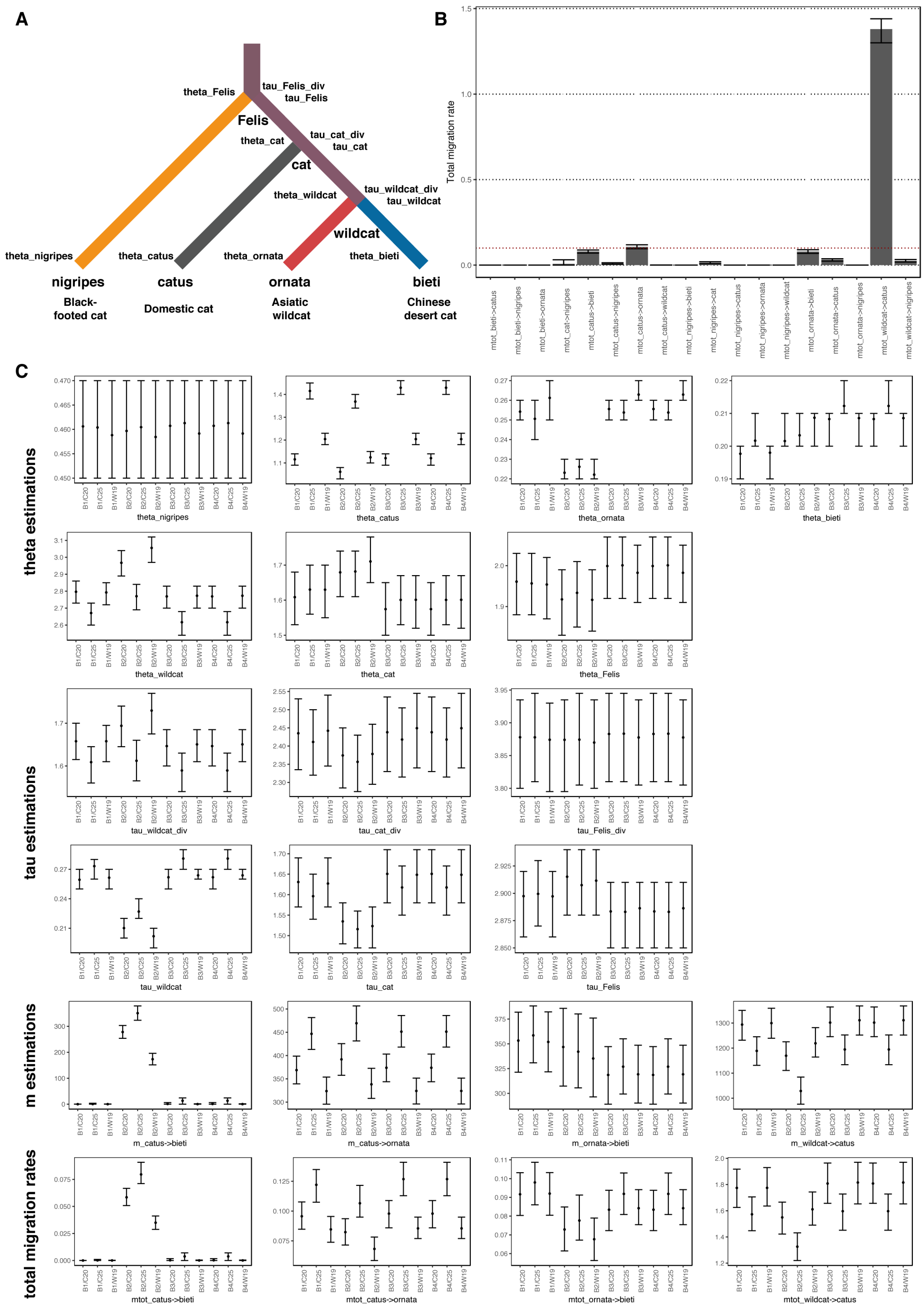

A
theta_Felis
tau_Felis_div
tau_Felis
Felis
theta_cat
tau_cat_div
tau_cat
cat
theta_wildcat
tau_wildcat_div
tau_wildcat
wildcat
theta_nigripes
theta_catus
theta_ornata
theta_bieti
nigripes
catus
ornata
bieti
Black-footed cat
Domestic cat
Asiatic wildcat
Chinese desert cat
B
Total migration rate
C
theta estimations
tau estimations
m estimations
total migration rates
theta_nigripes
theta_catus
theta_ornata
theta_bieti
theta_wildcat
theta_cat
theta_Felis
tau_wildcat_div
tau_cat_div
tau_Felis_div
tau_wildcat
tau_cat
tau_Felis
m_catus->bieti
m_catus->ornata
m_ornata->bieti
m_wildcat->catus
mtot_catus->bieti
mtot_catus->ornata
mtot_ornata->bieti
mtot_wildcat->catus

**Fig. S3.** Topology model and parameters estimated by G-PhoCS with four *F. s. bieti*, three *F. s. catus*, one *F. s. ornata* and one *F. nigripes*.

(A) Topology model and the parameters estimated by G-PhoCS analysis, with current and ancestral population number indicated below the nodes. Parameters related with Ne (theta) was marked at the left and parameters related with population divergence time (tau) and coalescent time was marked at the right.

(B) Total migration rates estimated in prior analysis with all the 18 possible migration bands included in the analysis. The height of the columns and error bars indicates average and the 95% HPD of estimations. Four migrations bands with a total migration rate over or around 0.1 (marked out with red dashed line) were identified as significant migration events and involved in final analysis.

(C) Estimations of theta, tau, m parameters and total migration rates in 12 independent analysis, with the Chinese mountain cat and domestic cat individuals involved in certain analysis noted on the X axis. The points and error bars indicate mean and 95% HPD intervals of the estimations respectively.

**Table S1.** Primers used in mitochondrial and Y-chromosome fragments amplification.

| **Fragment** | **Primer name** | **Sequence (5'-3')** | **Length (bp)** | **Reference** |
|---|---|---|---|---|
| Mitochondrial | CD-ND56-F1C | AACGTGGCTTTTTCAACTTTTA | 2686 | Driscoll et al. 2007 |
| | ND56-R2 | TCGGATGATTCAGCCATAAT | | |
| | CD-ND56-F1C | AACGTGGCTTTTTCAACTTTTA | 512 | |
| | ND56-1_R | TCTGCTCGGCCATATCATCA | | |
| | CD-ND56-F2B | TGCCGCCCTACAAGCAAT | 356 | |
| | CD-ND56-R4 | TTTGATATCATTTTGTGTGAGAGCAC | | |
| | ND56-1_F | TTATCCCCGTAGCGCTTTTC | 183 | This study |
| | ND56-1_R | TCTGCTCGGCCATATCATCA | | |
| | ND56-2_F | CCCCAACAACTATCCCACAA | 158 | |
| | ND56-2_R | TTGATGGGAGGCGGTGTATT | | |
| Y chromosome | DBY7_F | GGTCCAGGAGARGCTTTGAA | 317 | Hellborg et al. 2003 |
| | DBY7_R | CAGCCAATTCTCTTGTTGGG | | |
| | SMCY7_F | TGGAGGTGCCCRAARTGTA | 698 | |
| | SMCY7_R | AACTCTGCAAASTRTACTCCT | | |

**Table S2.** Pairwise $F_{ST}$ between different groups estimated based on 1Mb windows on autosomes

| | ***F. nigripes*** | ***F. s. bieti*** | ***F. s. ornata*** | ***F. s. catus* (Chinese)** | ***F. s. catus* (Worldwide)** |
|---|---|---|---|---|---|
| ***F. nigripes*** | - | 0.037 | 0.029 | 0.051 | 0.051 |
| ***F. s. bieti*** | 0.935* | - | 0.121 | 0.069 | 0.059 |
| ***F. s. ornata*** | 0.950 | 0.785 | - | 0.112 | 0.097 |
| ***F. s. catus* (Chinese)** | 0.736 | 0.504 | 0.413 | - | 0.040 |
| ***F.s. catus* (Worldwide)** | 0.691 | 0.443 | 0.297 | 0.104 | - |

* Numbers below and above the diagonal are averages and standard errors of the estimations

**Table S3.** Genetic introgression of *F. s. bieti* into *F. s. catus* from different areas of China.

| | ABBA-BABA test | | Diagnostic sites | *f*-statistics | | *f*4-ratio test | |
|---|---|---|---|---|---|---|---|
| WGS code | Average D-stat | Average Z score | Percentage (%) | S-stat | Percentage (%) | Percentage (%) | Z score |
| C1 | 0.007 | 1.25 | 1.63 | 3750 | 0.20 | 3.32 | 3.10 |
| C2 | 0.010 | 2.03 | 1.61 | 6530 | 0.35 | 7.51 | 5.17 |
| C3 | 0.015 | 2.23 | 1.95 | 10084 | 0.53 | 7.18 | 5.13 |
| C4 | 0.023 | 3.17 | 2.71 | 17192 | 0.91 | 4.93 | 5.06 |
| C5 | 0.012 | 1.87 | 2.11 | 7524 | 0.40 | 2.91 | 3.83 |
| C6 | 0.202 | 12.69 | 11.35 | 189489 | 10.03 | 11.20 | 8.93 |
| C7 | 0.205 | 13.41 | 11.43 | 194344 | 10.28 | 10.85 | 9.03 |
| C8 | 0.157 | 9.25 | 8.65 | 142353 | 7.53 | 9.30 | 6.44 |
| C9 | 0.210 | 13.67 | 11.64 | 198688 | 10.51 | 11.31 | 9.85 |
| C10 | 0.138 | 10.09 | 7.79 | 123393 | 6.53 | 8.20 | 7.50 |
| C11 | 0.076 | 6.94 | 4.99 | 63856 | 3.38 | 4.88 | 5.31 |
| C12 | 0.193 | 12.79 | 10.70 | 180314 | 9.54 | 10.42 | 9.24 |
| C13 | 0.105 | 7.95 | 6.22 | 90812 | 4.81 | 7.16 | 6.66 |
| C14 | 0.105 | 8.52 | 6.25 | 92220 | 4.88 | 6.99 | 6.28 |
| C15 | 0.075 | 6.99 | 4.88 | 62598 | 3.31 | 4.45 | 4.59 |
| C16 | -0.020 | -4.60 | 0.81 | -18667 | -0.99 | -2.05 | -2.35 |
| C17 | -0.018 | -3.67 | 0.94 | -17055 | -0.90 | -1.96 | -2.09 |
| C18 | -0.014 | -2.98 | 1.19 | -13981 | -0.74 | -0.04 | -0.04 |
| C19 | -0.018 | -4.20 | 0.99 | -16745 | -0.89 | -3.03 | -3.91 |
| C20 | -0.026 | -6.03 | 0.63 | -23628 | -1.25 | -2.99 | -3.22 |
| C21 | -0.028 | -6.71 | 0.70 | -25258 | -1.34 | -3.32 | -3.94 |
| C22 | -0.020 | -4.74 | 0.78 | -18639 | -0.99 | -0.79 | -0.89 |
| C23 | -0.021 | -4.67 | 0.91 | -19400 | -1.03 | -1.83 | -1.94 |
| C24 | -0.023 | -5.38 | 0.85 | -21173 | -1.12 | -1.18 | -1.22 |
| C25 | -0.023 | -5.47 | 0.89 | -20972 | -1.11 | -1.71 | -1.94 |
| C26 | -0.018 | -4.13 | 0.80 | -16951 | -0.90 | -2.80 | -3.28 |
| C27 | -0.022 | -5.08 | 0.84 | -20610 | -1.09 | -2.29 | -2.53 |
| C28 | -0.022 | -5.05 | 0.91 | -20075 | -1.06 | -2.43 | -2.63 |
| C29 | -0.016 | -3.52 | 0.73 | -14922 | -0.79 | -1.17 | -1.25 |
| C30 | -0.014 | -3.15 | 0.74 | -13227 | -0.70 | -2.98 | -3.94 |
| C31 | -0.018 | -4.41 | 0.65 | -17086 | -0.90 | -2.30 | -2.93 |
| C32 | -0.022 | -5.52 | 0.58 | -20165 | -1.07 | -3.59 | -4.91 |
| C33 | -0.023 | -5.36 | 0.74 | -20565 | -1.09 | -3.52 | -4.43 |
| C34 | -0.023 | -5.30 | 0.70 | -20969 | -1.11 | -2.60 | -2.89 |
| C35 | -0.021 | -4.92 | 0.61 | -18978 | -1.00 | -3.43 | -4.23 |
| C36 | -0.015 | -3.28 | 0.71 | -14458 | -0.77 | -1.82 | -2.05 |
| C37 | -0.027 | -6.36 | 0.63 | -23939 | -1.27 | -3.31 | -4.17 |
| C38 | -0.035 | -9.17 | 0.53 | -30281 | -1.60 | -5.30 | -7.23 |
| C39 | -0.022 | -5.32 | 0.70 | -20295 | -1.07 | -3.10 | -3.67 |
| C40 | -0.026 | -6.35 | 0.60 | -23034 | -1.22 | -4.80 | -6.71 |
| C41 | -0.028 | -7.26 | 0.54 | -24667 | -1.31 | -5.20 | -7.58 |
| C42 | -0.028 | -6.95 | 0.56 | -24714 | -1.31 | -4.25 | -5.91 |
| C43 | -0.031 | -7.56 | 0.57 | -26946 | -1.43 | -4.50 | -6.02 |
| C44 | -0.028 | -6.98 | 0.68 | -25026 | -1.32 | -4.67 | -6.38 |
| C45 | -0.030 | -7.65 | 0.70 | -26979 | -1.43 | -4.74 | -5.97 |
| C46 | -0.028 | -6.88 | 0.57 | -25047 | -1.33 | -4.35 | -5.23 |

**Table S4.** Alder fitting statistics with different fit starts estimated from weighted LD curves of two hybrid domestic cat populations.

| Pop | Fit start d0 (cM) | p-value | Number of generations | Amplitude of exponential | Z score |
|---|---|---|---|---|---|
| hybrid1 | | | | | |
| | 0.5 | 7.90E-15 | 8.07 +/- 1.04 | 0.00111125 +/- 0.00006854 | 7.77 |
| | 1.0 | 2.20E-15 | 7.77 +/- 0.98 | 0.00107075 +/- 0.00006901 | 7.93 |
| | 1.5 | 1.00E-15 | 7.57 +/- 0.94 | 0.00104127 +/- 0.00006998 | 8.02 |
| | 2.0 | 8.30E-16 | 7.42 +/- 0.92 | 0.00101923 +/- 0.00007108 | 8.05 |
| | 2.5 | 1.10E-15 | 7.30 +/- 0.91 | 0.00100081 +/- 0.00007256 | 8.02 |
| | 3.0 | 1.50E-15 | 7.20 +/- 0.90 | 0.00098502 +/- 0.00007412 | 7.98 |
| | 3.5 | 2.80E-15 | 7.12 +/- 0.90 | 0.00097259 +/- 0.00007603 | 7.9 |
| | 4.0 | 6.10E-15 | 7.06 +/- 0.90 | 0.00096177 +/- 0.00007841 | 7.8 |
| | 4.5 | 1.60E-14 | 7.00 +/- 0.91 | 0.00095246 +/- 0.00008105 | 7.68 |
| | 5.0 | 4.90E-14 | 6.97 +/- 0.92 | 0.00094591 +/- 0.00008397 | 7.53 |
| hybrid2 | | | | | |
| | 0.5 | 8.90E-05 | 30.72 +/- 7.84 | 0.00022193 +/- 0.00002941 | 3.92 |
| | 1.0 | 0.00033 | 22.94 +/- 6.40 | 0.00016569 +/- 0.00003023 | 3.59 |
| | 1.5 | 0.00079 | 18.83 +/- 5.61 | 0.00013311 +/- 0.00002649 | 3.36 |
| | 2.0 | 0.0021 | 16.50 +/- 5.36 | 0.00011394 +/- 0.00002399 | 3.08 |
| | 2.5 | 0.0065 | 14.99 +/- 5.51 | 0.00010128 +/- 0.00002307 | 2.72 |
| | 3.0 | 0.018 | 14.21 +/- 6.01 | 0.00009461 +/- 0.00002450 | 2.36 |
| | 3.5 | 0.039 | 13.74 +/- 6.65 | 0.00009060 +/- 0.00002779 | 2.07 |
| | 4.0 | 0.073 | 13.46 +/- 7.50 | 0.00008808 +/- 0.00003345 | 1.8 |
| | 4.5 | 0.14 | 12.88 +/- 8.66 | 0.00008297 +/- 0.00004200 | 1.49 |
| | 5.0 | 0.22 | 12.38 +/- 10.00 | 0.00007857 +/- 0.00005185 | 1.24 |

**Table S5.** Summary statistics of G-PhoCS results, with average and combined 95% HPD of parameters from different runs and effective populations sizes, divergence times and total migration rates calculated with these estimated parameters.

| | theta (E-03) | | | Effective population size (E+04) | | |
|---|---|---|---|---|---|---|
| **Statistics** | **mean** | **combined 95% HPD** | | **mean** | **combined 95% HPD** | |
| theta_nigripes | 0.46 | 0.45 | 0.47 | 4.42 | 4.33 | 4.52 |
| theta_bieti | 0.21 | 0.19 | 0.22 | 2.02 | 1.83 | 2.12 |
| theta_ornata | 0.24 | 0.22 | 0.27 | 2.31 | 2.12 | 2.60 |
| theta_catus | 1.23 | 1.03 | 1.46 | 11.83 | 9.90 | 14.04 |
| theta_wildcat | 2.78 | 2.54 | 3.12 | 26.73 | 24.42 | 30.00 |
| theta_cat | 1.62 | 1.50 | 1.78 | 15.58 | 14.42 | 17.12 |
| theta_Felis | 1.97 | 1.83 | 2.07 | 18.94 | 17.60 | 19.90 |
| | **tau (E-03)** | | | **Divergence time (Mya)** | | |
| tau_wildcat | 0.25 | 0.19 | 0.29 | 0.19 | 0.15 | 0.22 |
| tau_cat | 1.61 | 1.47 | 1.71 | 1.24 | 1.13 | 1.32 |
| tau_Felis | 2.89 | 2.85 | 2.94 | 2.22 | 2.19 | 2.26 |
| tau_wildcat_div | 1.64 | 1.54 | 1.77 | 1.26 | 1.18 | 1.36 |
| tau_cat_div | 2.42 | 2.28 | 2.55 | 1.86 | 1.75 | 1.96 |
| tau_Felis_div | 3.88 | 3.80 | 3.95 | 2.98 | 2.92 | 3.04 |
| tau_div | 3.88 | 3.88 | 3.88 | 2.00 | 3.00 | 3.00 |
| | **m** | | | **Total migration rate** | | |
| m_ornata->bieti | 334.80 | 289.08 | 388.25 | 0.09 | 0.06 | 0.11 |
| m_catus->ornata | 386.46 | 295.54 | 506.62 | 0.10 | 0.06 | 0.14 |
| m_catus->bieti | 69.36 | 0.00 | 378.45 | 0.02 | 0.00 | 0.09 |
| m_wildcat->catus | 1234.55 | 975.96 | 1368.21 | 1.67 | 1.22 | 1.97 |

**Data file S1.** Wildcat and domestic cat samples collected in the study, including the sample information and genetic information from preliminary analysis.

**Data file S2.** Haplotypes and variable sites identified in the 2.6 kb mtDNA fragment.

**Data file S3.** Haplotypes and variable sites identified in the 1016 bp concatenated Y-chromosome fragment.

**Data file S4.** Statistics of WGS reads generated and mapped to felCat8 assembly, domestic cat Y-chromosome and mitochondrial genome sequence.
